# A multilayer network approach elucidates time- and tissue-specific developmental and aging processes

**DOI:** 10.1101/2025.05.02.651926

**Authors:** Salvo D. Lombardo, André F. Rendeiro, Jörg Menche

## Abstract

Understanding the dynamic interplay of proteins across different life stages and tissues is essential for deciphering the molecular mechanisms underpinning development, aging, and disease. Here, we present a comprehensive network-based framework that constructs and integrates 119 time- and tissue-specific protein-protein interaction (PPI) networks derived from transcriptomic data, offering insights into proteomic dynamics across the human lifespan. Based on this, we observed three distinct protein groups: (i) *common-core proteins*, expressed universally across all tissues and time points; (ii) *time-/tissue-specific proteins*, selectively expressed within specific temporal or spatial contexts; and (iii) *time-/tissue-unique proteins*, whose expression is restricted to specific points in space and time. Our analysis shows a clear gradient of network centrality, transitioning from the highly connected common-core proteins to more specialized time-/tissue-specific and unique proteins, mirroring a progressive shift in functional specificity. Further, we characterized the distinct molecular signatures of intrauterine to extrauterine life, delineating two key protein networks: the *embryonic development network (EDev)* and the *environmental aging network (EAgi)*. Their network characterization and comparison highlighted specific communities within the EDev network enriched for developmental diseases, and specific EAgi communities involved in aging. This network classification allowed us to rank candidate anti-aging drugs and their molecular targets, laying the foundation for a systematic, data-driven, network-based investigation of development and aging, providing a roadmap for future research aimed at mitigating age-related diseases and promoting longevity.

## Introduction

Aging and development are critical time-dependent processes, yet their underlying molecular mechanisms remain only partially understood. Establishing parallelisms between these phenomena may help identify shared and unique molecular pathways. For example, recent studies have suggested that the same signaling pathways involved in development also play a role in regulating health span and aging ^1^, with their dysregulations contributing to diseases such as cancer, dementia, and developmental malformations ^2^. While this idea is compelling, a systematic molecular comparison of these two biological processes is still lacking. The extensive availability of transcriptomic studies over the past decades provides an opportunity for a data-driven, unbiased, systematic overview of the molecular processes underpinning development ^3^ and aging ^4^. However, since proteins do not act in isolation, it is essential to consider their broader context by mapping their interactions and the protein complexes they form. Protein-protein interaction (PPI) networks, or interactomes, provide a framework for such analyses. These versatile tools have been successfully applied to reveal how groups of interacting proteins lead to disease phenotypes ^5,6^. To model aging and development, defining context-specific PPI networks is crucial to capture interactions unique to specific tissues or life stages. For example, studies have shown that many Mendelian diseases display tissue-specific phenotypes that are not fully explained by tissue expression profile alone, but can be understood by focusing on tissue-specific PPIs ^7^. However, elucidating both tissue- and temporal-specificity in PPIs remains unexplored.

In this study, we propose a framework to construct time- and tissue-specific PPI networks derived from transcriptomic data. Using this approach, we generated 119 networks with nodes representing highly expressed proteins and their immediate interactors within the global human PPI network. The overlap of these networks highlighted a core set of 2371 proteins consistently expressed across all life stages, which are highly connected in the global human PPI network. These proteins were found to be enriched for housekeeping functions, forming densely connected protein complexes. To investigate molecular differences between intra- and extra-uterine life, we grouped the time- and tissue-specific PPI networks accordingly. While many proteins were shared between these two periods, our analyses uncovered key differences in network connectivity, which may contribute to the differences observed between development and aging. We further characterized these processes, identifying network densely connected molecular mechanisms unique to development, aging or shared by both, that we have described through various biologically enrichment analyses. Finally, using network-based distances, we also identified candidate anti-aging compounds targeting these molecular pathways, suggesting potential therapeutic interventions to modulate aging-related processes.

## Results

### Definition of the multi-layer time/tissue-specific network

To decipher the intricate transcriptomic changes occurring in several tissues in different periods of life, we interrogated data from ^8^, which comprehends bulk RNA-seq expression for seven distinct tissues over time. This dataset provides a panoramic view of the developmental trajectories spanning the three embryonic germ layers: ectoderm (comprising brain and cerebellum), mesoderm (encompassing heart, kidney, ovary, and testis), and endoderm (represented by the liver). The temporal resolution of this dataset spans from the 4th-week post-conception (WPC) to 65 years of age, allowing us to trace tissue-specific trajectories from development through various life stages, thereby characterizing their aging processes.

We derived tissue- and time-specific PPI networks through a three-step methodology. Initially, we identified the most highly expressed genes within each of the seven tissues using the Genotype-Tissue Expression (GTEx) dataset. Subsequently, we computed a discerning cut-off to select highly expressed corresponding transcripts at each temporal point for every tissue (see methods). These have been mapped into a comprehensive PPI network characterized by 18.815 nodes and 482.935 edges ^9^ and used as seed proteins for detecting their immediate neighbors through a random walk with restart algorithm (Fig. 1A). Ultimately, we obtained 119 PPI networks, whose number of nodes ranged from 5,589 to 10,626 and whose number of edges from 194,010 to 408,146. Notably, we observed a decreasing trend in the time- and tissue-specific expression TPM cutoff over time (Supplementary Fig. 1A), which corresponded to a reduction in the number of seed proteins identified (Supplementary Fig. 1B).This observation aligns with previous studies reporting that aging is associated with an overall reduced activity in growth factors, mitochondrial function, and protein synthesis ^10^. Despite this trend, the overall number of nodes and edges in the final networks was not affected over time (Supplementary Fig. 1C, D). Interestingly, the TPM cutoff was not statistically significantly associated with the number of nodes but with the number of edges of each network

**Figure 1:**
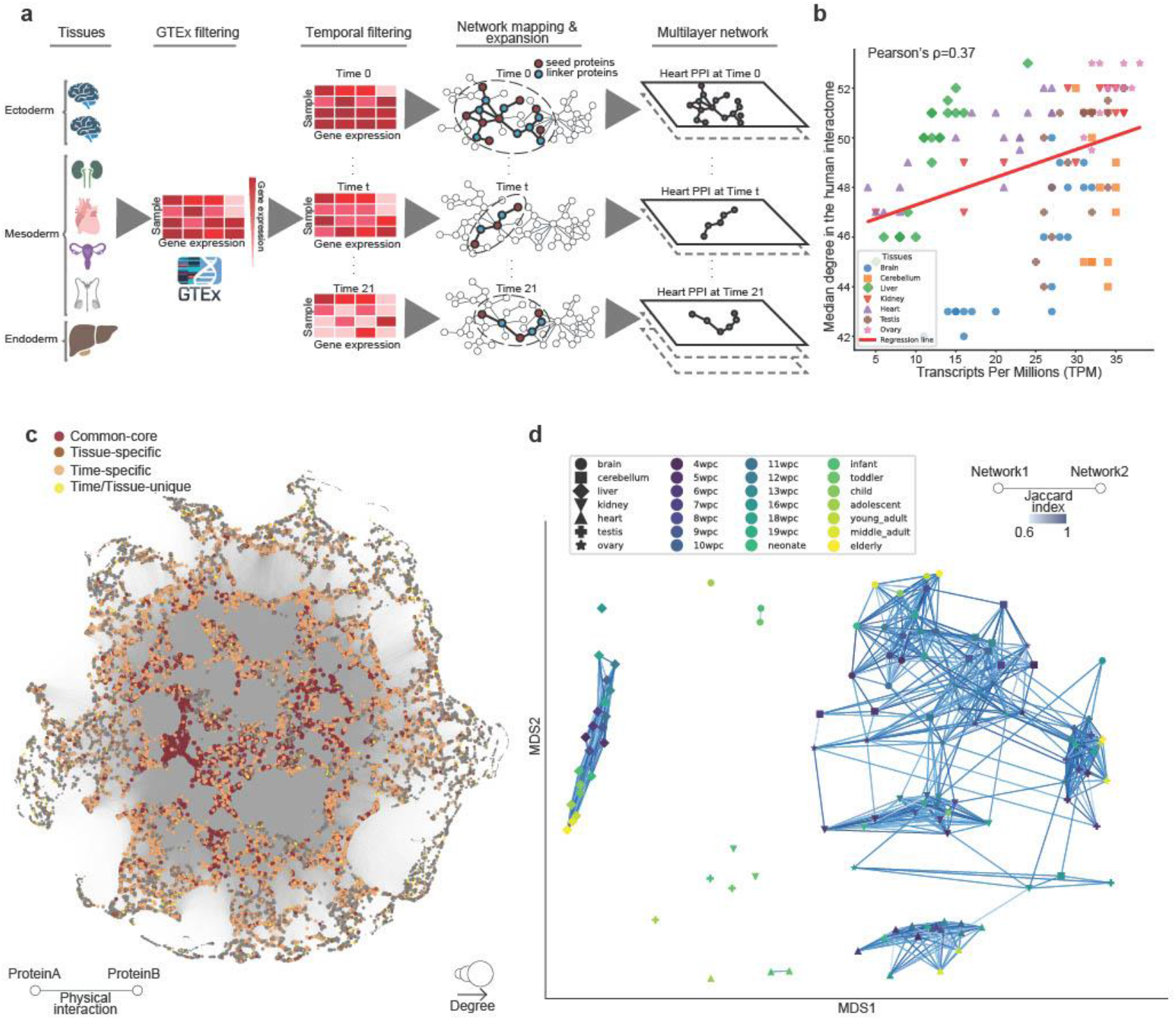
Definition of Time- and Tissue- specific multilayer PPI network. **A**, Illustrative process of the creation of the time- and tissue- specific multilayer PPI network. The transcriptomic temporal dynamics of three embryonic germ layers is captured by the RNA-seq expression of 7 distinct tissues: brain, cerebellum, kidney, heart, ovary, testis, and liver. Two expression cut-offs were applied, a tissue-specific cutoff, which was calculated by using GTEx as a reference, and a temporal cutoff, which is based on Cardoso-Moreira tissue expression. Finally, this set of highly tissue- and time-specific proteins was expanded by a random walk with restart in the PPI, until the largest connected component that included all seed proteins was reached. **B**, Scatterplot showing the correlation between the median expression of the seed proteins for each time-/tissue-specific network (expressed in TPM) and the median degree in the global human interactome (protein-protein interaction network). **C**, Visualization of the global human interactome, each node represents a protein, and an edge a physical interaction. The size of each node is proportional to its own degree, while the color reflects the property of that protein: red (common-core protein, expressed at all times and tissues), brown (tissue-specific protein, expressed at all times in a specific tissue, but excluding common-core proteins), orange (time-specific protein, expressed in every tissue for a given time, and excluding common-core proteins), yellow (time/tissue unique proteins, expressed only in a specific tissue in a given time). **D**, Multidimensional scaling plot showing how networks derived from the same tissue or closer in time tend to cluster together. Each dot represents a tissue-/time-specific PPI network, each tissue is represented by a different shape, the gradient color follows the temporal evolution (from dark blue to light yellow). An edge is drawn between two networks, if they share at least 60% of their edges and their width is proportional to the edge overlap between the two networks.

(Pearson’s *ρ*=0.2, *p-value*=0.03, Supplementary Fig. 2D), suggesting that more stringent TPM cutoffs may lead to more central proteins in the human PPI. To validate this hypothesis, we assessed the relationship between the TPM cutoff and the median degree of seed proteins in the human PPI network for each tissue and time combination. We found a positive correlation between these variables (Pearson’s *ρ*=0.37, *p-value* = 2e-5, Fig. 1B), indicating that higher stringency in expression cutoffs indeed enriches for proteins with greater centrality in the interactome.

Besides being robust on the initial number of seed genes, our approach is capable of capturing tissue- and time-variability across the 119 time-/tissue-specific networks, as shown by the broad range of edge overlap values (0.45-0.97), calculated using the Jaccard Index (JI) (Supplementary Fig. 3A), with a higher similarity for those comparisons involving networks originating from the same tissue or derived from adjacent times (Supplementary Fig. 3B, Supplementary Table 1A). Intriguingly, the liver displays the fewest number of conserved edges over time (163,230 edges), while the ovary the highest (258,963 edges) (Supplementary Fig. 3C). Notably, all tissue comparisons showed a statistically significant number of shared edges (Fisher’s exact test, Supplementary Table 1B; Supplementary Fig. 3D). To refine our analysis and pinpoint tissue-specific pathways, we focused on overlapping proteins that formed a statistically significant largest connected component within their respective networks (i.e., participating in the same biological machinery, see methods). However, even with this refinement, the average number of shared proteins remained high (4787 across all tissue comparisons), obscuring tissue-specific pathways (Supplementary Table 1B). Similar trends were observed in temporal comparisons (Supplementary Table 1C; Supplementary Fig. 3E, F).

To address these limitations, we treated the 119 PPI networks as layers of a multi-layer network, enabling a systematic integration of both tissue and time components. Through this approach, we found that over 50% of all edges were shared across all layers, identifying a core set of 2371 proteins persistently present across all tissues and time points (“*common-core*”). We then distinguished *tissue-specific* and *time-specific proteins*, which are those present in the same time- and tissue-layers respectively (but excluding the common-core proteins), and finally, the *time/tissue unique proteins* that are present only in a specific layer of the multi-layer network (i.e. in a precise time and tissue) (Fig. 1C). This new classification allowed us to enhance the observed variability across all pairwise network comparisons, highlighting differences and commonalities between tissues (JI_min_= 0.14 (between liver and cerebellum), JI_max_= 0.63 (between brain and cerebellum)) (Fig. 1D; Supplementary Fig. 4A; Supplementary Table 2A), and times (Supplementary Fig. 4B; Supplementary Table 2B). Furthermore, the average number of shared proteins participating in significantly connected pathways was reduced by 4-fold, allowing for clearer identification of those molecular pathways enriched in specific *tissue* ***X*** *times* combination (Supplementary Table 2A, B). By focusing on the *time- and tissue-specific proteins*, we were able to elucidate differences and similarities between tissues more effectively across all observed time points, as shown in the multidimensional scaling plot in Fig. 1D, where clear tissue clustering emerged, alongside a temporal gradient within each tissue (Fig. 1D; Supplementary Table 2C).

### Topological and biological characterization of the common-core proteins

The examination of the 2371 common-core proteins revealed cohesive interconnectivity among them, observing a large divergence on their largest connected component compared to a 1,000 random protein sets of the same size (*z-score* = 14.5, Supplementary Fig. 5A). Predominantly, these proteins served as seed in at least one network layer (95%, n=2250), with 45% (n=1070) acting as seed proteins in all networks, which highlights their widespread high expression across tissues. Based on these characteristics, we hypothesize their involvement in broad biological activities. To test this, we used network topological considerations whose findings have been validated across several approaches and data modalities presented below.

Numerous studies have shown a robust correlation between network centrality in the interactome and biological activity ^11,12^. Along this line, we postulated that common-core proteins exhibit higher centrality in our tissue-time-specific interactomes; on the contrary, proteins with more specific biological roles would be more peripheral in the corresponding PPI network. Our analyses support this hypothesis, with common-core proteins having the highest degree, while tissue/time-unique proteins showing the smallest (Fig. 2A). Interestingly, while the number of common-core proteins remained constant across all networks (by definition), the number of time-specific proteins showed distinct peaks at specific periods in life: 8wpc, young adulthood, and elderly (Supplementary Fig. 5B). Since the definition of time-specificity depends on the overlap of proteins expressed within networks for the same period, it is indirectly influenced by the number of available samples. To test this, we examined the relationship between the number of networks available at each time point and the number of time-specific proteins. Although a negative correlation was observed, it was not statistically significant (Pearson’s *ρ*=-0.36, *p-value*:0.1, Supplementary Fig. 5C). To gain deeper biological insights into these time-points, we focused on those time-specific proteins appearing for the first time at a given time point, referring to them as *‘unique’ time-specific proteins*. While the number of unique time-specific proteins decreases over time (by definition), the same peaks were observed at 8wpc, young adulthood and elderly (Supplementary Fig. 6A). Counterintuitively, the number of unique time-specific proteins showed a weak, non-significant, positive correlation with the number of networks available for each time-point (Pearson’s *ρ*=0.22, *p-value*:0.32, Supplementary Fig. 6B). This trend can be attributed to the higher availability of samples in earlier time points (intrauterine life), which is inherently enriched with a larger number of unique time-specific proteins. Furthermore, we found that the network centrality of unique time-specific proteins generally decreased over time, except at the same three peaks (8wpc, young adulthood, and elderly) (Supplementary Fig. 6D). From a biological perspective, these unique time-specific proteins explicate distinct, timely regulated functions. For instance, at 8wpc they were enriched in biological processes related to vesicle-mediated transport (*adjusted p-value*: 0.04, Supplementary Fig. 7A), and cellular components of the clathrin system (*adjusted p-value*: 0.01, Supplementary Fig. 7C), suggesting a role in the fetal DNA delivery system. Remarkably, this timing aligns with the 8th week post-conception (10th week of pregnancy), point at which fetal DNA becomes detectable in maternal blood through non-invasive prenatal testing (NIPT) ^13^.

**Figure 2:**
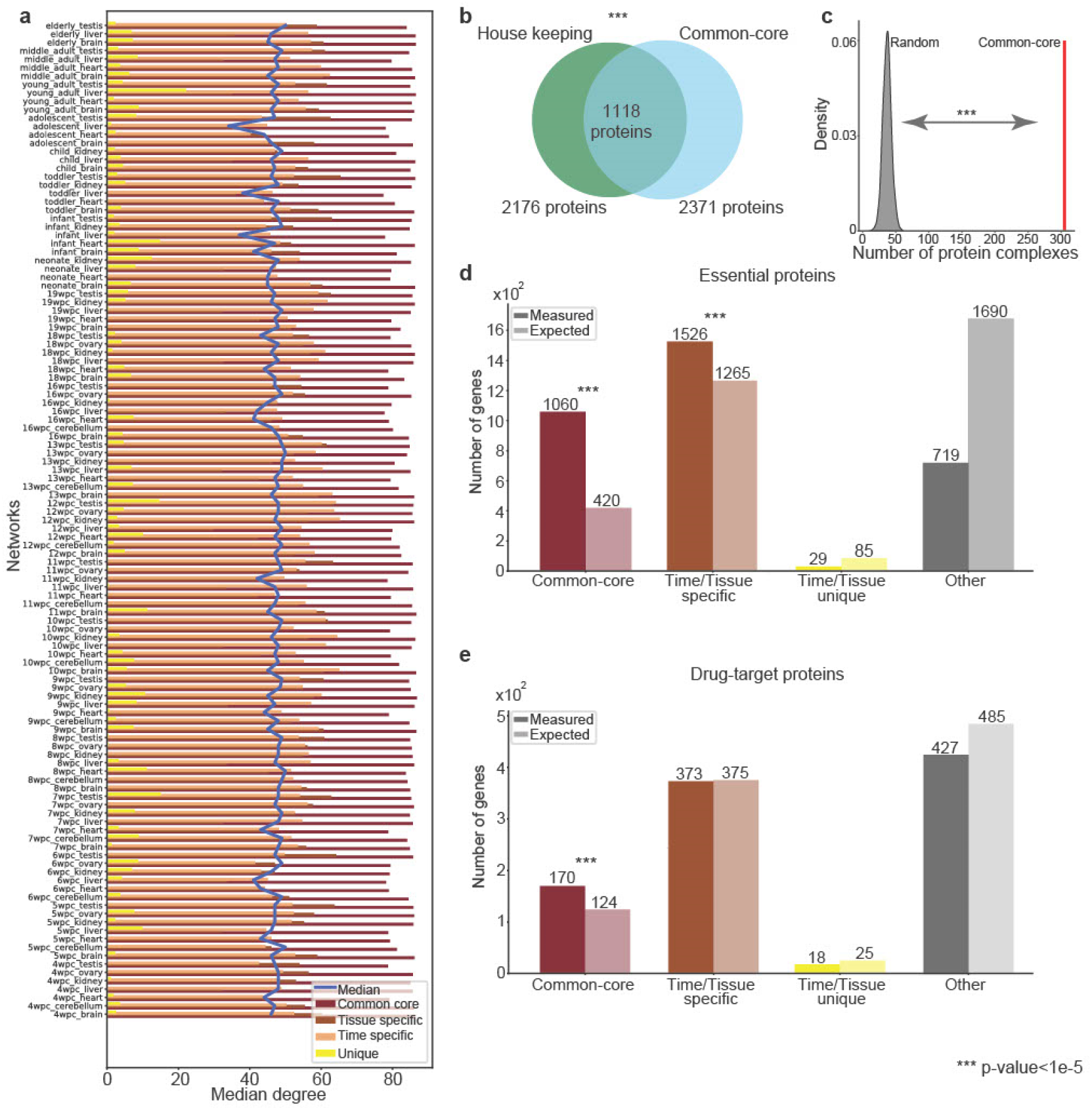
Common-core proteins are crucial in human biology. **A**, Barplot showing the median degree on all tissue/time-specific networks of different groups of proteins: common-core (dark-red), tissue-specific (dark-brown), time-specific (light-brown), unique (yellow). A blue line shows the overall median degree across all tissue/time-specific networks. **B**, Venn-diagram showing the overlap between common-core proteins and Housekeeping proteins. **C**, Distribution of the number of protein complexes selecting an equal number of random proteins to the common core for 10,000 times (z-score = 41). **D**, Barplot showing the number of measured essential proteins in different groups (common-core, time/tissue-specific, time/tissue-unique, others) compared to their expectation. **E**, Barplot showing the number of measured drug-target proteins in different groups (common-core, time/tissue-specific, time/tissue-unique, others) compared to their expectation.

In contrast, the biological enrichment of the common-core proteins was indicative of fundamental cellular functions occurring at all life stages. Specifically, 47% of these proteins were identified as housekeeping proteins (*p-value*<1e-299, Fisher’s exact test, Fig. 2B), highlighting their essential role in maintaining basic cellular activities. Moreover, these proteins tend to participate in a larger number of protein complexes than expected by random chance (*z-score* = 41, Fig. 2C), underscoring their involvement in intricate molecular interactions crucial for cellular processes. Furthermore, we observed that essential genes are not equally distributed across the 4 categories we defined (*p-value*: 2e-299, multinomial test, Fig. 2D), but instead they enriched with both common-core proteins (*p-value*: 9e-142, Fisher’s exact test), and time/tissue-specific proteins (*p-value*: 9e-32, Fisher’s exact test), but not with tissue/time unique proteins or unclassified proteins. Similarly, the drug-target proteins of the 908 FDA-approved drugs reported in DrugBank enriched with the common-core (*p-value*: 1e-05, Fisher’s exact test; *p-value*: 4e-09, multinomial test, Fig. 2E), confirming the relevance of these proteins from a therapeutic point of view. Exploring the disease phenotype, we found that proteins involved in Autosomal Dominant Mendelian diseases are enriched with common-core proteins (*p-value*: 2e-13, Fisher’s exact test), but not the Autosomal Recessive ones, which instead enriched for time/tissue-specific proteins (*p-value*: 4e-7, Fisher’s exact test), suggesting a milder biological impact of time/tissue-specific proteins compared to the common-core ones (Supplementary Fig. 7D).

Finally, we compared our findings with an external dataset (BBI) ^14^ comprising single-cell RNA-seq data during development for five of the seven analyzed tissues (brain, cerebellum, heart, kidney, and liver). We observed a positive correlation of the normalized transcriptomic expression (transcripts per kilobase million, TPM) between the two datasets (average Spearman’s *ρ*: 0.74, Supplementary Fig. 8A). Furthermore, while highly expressed proteins in each tissue from BBI tend to enrich with common-core proteins and time/tissue-specific proteins (Supplementary Fig. 8B; Supplementary Table 3A), the highly frequent proteins tend to enrich in time/tissue-specific proteins of the corresponding tissue (Supplementary Fig. 8C; Supplementary Table 3B), and lastly, the differentially expressed proteins of the most common cell type for each tissue tend to be highly represented in the tissue/time-specific and unique proteins (Supplementary Fig. 8D; Supplementary Table 3C). These consistent trends across datasets underscore the robustness and biological relevance of our observations, reinforcing the validity of our initial hypothesis that common-core proteins are involved in a broad range of biological activities.

However, because proteins don’t act in isolation but rather exert their function through physical interactions, our protein classification was extended to the edge level, encompassing 10 distinct types of protein-protein interactions (Supplementary Fig. 9A). As expected, the vast majority of edges in the PPI network involve at least one common-core protein (73%), despite these proteins representing only 12.7% of all proteins in the PPI network. This observation aligns with the high degree exhibited by common-core proteins in the PPI. On the contrary, unclassified proteins represent 47% of the total PPI, yet the interactions among them are limited (1.5% of all edges in the PPI), and the fraction of edges in which an unclassified protein takes part is about 20% of all potential interactions (Supplementary Fig. 9B). Intrigued by this disproportion, we compared the network density of each of these groups of proteins against the overall density of the PPI, revealing the highest density for interactions involving two common-core proteins (CC) (Supplementary Fig. 9C). In a scenario where network density is uniformly distributed across all groups, we would expect 7,593 edges in the CC group, and 105,063 edges in the OO (other-to-other) group. Instead, we observed 126,002 and 7,106 respectively. This discrepancy holds significant biological implications. Despite having the fewest number of proteins, the highest number of statistically significant enriched terms (*FDR*<0.05, Fisher’s exact test with Benjamini-Hochberg correction) for the Gene Ontology (GO) was found with interactions that involve at least one common-core protein (Supplementary Fig. 9D; Supplementary Fig. 10; Supplementary Table 4). Notably, common-core proteins are responsible for essential biological activities including protein transcription and translation, operating in both the nucleus and cytoplasm compartments.

### Definition of the embryological developmental network

Our next goal was to distinguish developmental-specific molecular mechanisms from those that are influenced by environmental factors. To address this, we grouped all networks into two classes (before and after birth) and evaluated whether the presence of time-specific proteins could predict their temporal period. To achieve this, we applied three distinct machine learning (ML) classifiers: logistic regression, support vector classifier (SVC), and random forest. Given the limited dataset size (number of networks = 119), we employed a “leave-one-out” cross-validation strategy. This approach yielded an average accuracy of approximately 70% in matching the predicted temporal period with the ground truth with all three methods (Fig. 3A; Supplementary Fig. 11; Supplementary Table 5).

**Figure 3:**
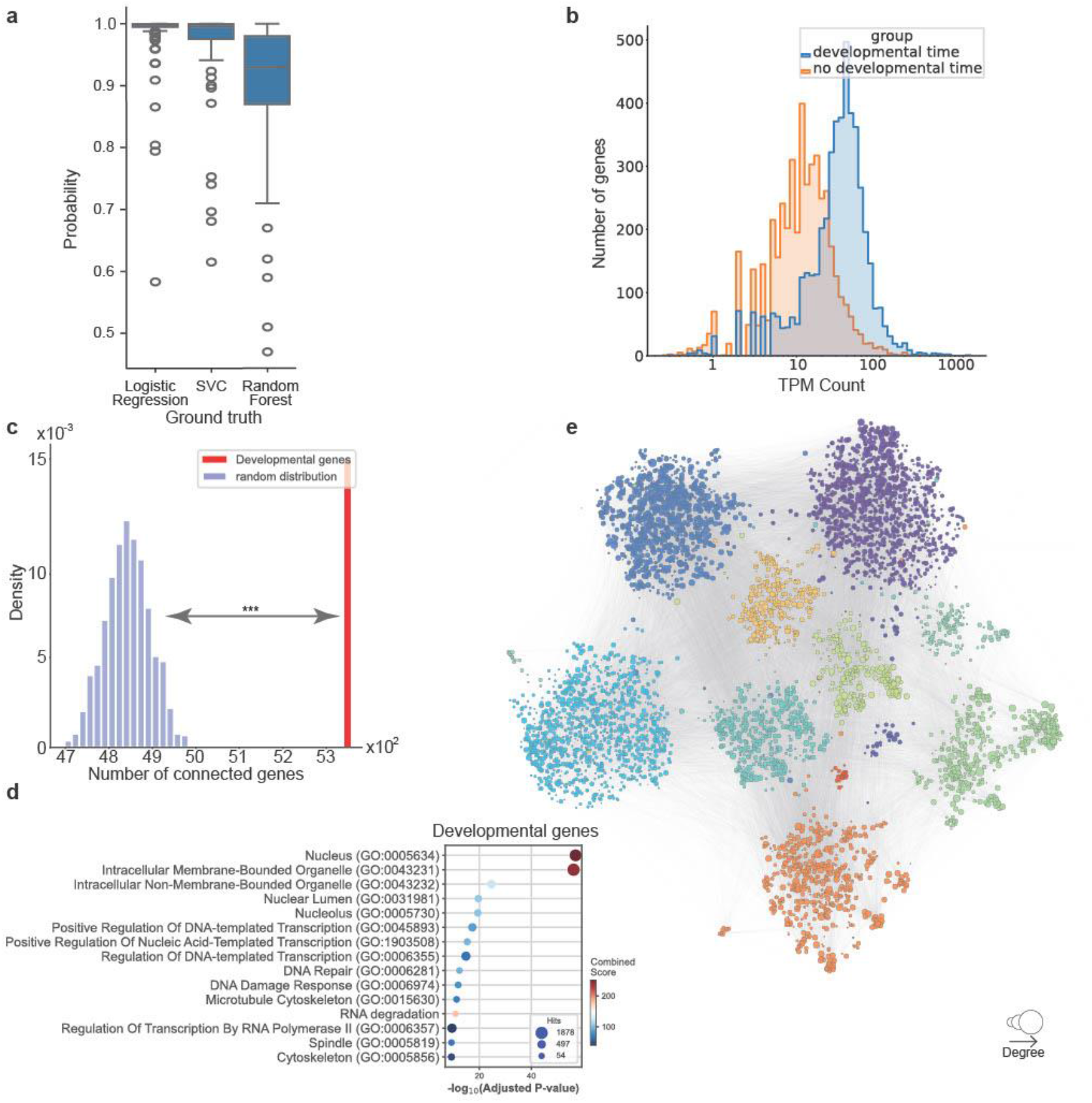
Intrauterine-specific proteins form an interconnected embryological developmental (EDev) network. **A**, Boxplot showing the accuracy to classify each time-/tissue-specific network in its proper time period. Three algorithms have been used: Logistic Regression, Support Vector Classifier (SVC) and Random Forest. **B**, Transcriptomic expression comparison of developmental and no-developmental time-specific proteins. **C**, Number of connected proteins in the largest connected component of the developmental time-specific proteins. **D**, Statistically significant enriched terms of the developmental time-specific proteins (FDR<0.05) with the three branches of the protein Ontology: Biological Processes (BP), Molecular Functions (MF), Cellular Components (CC). **E**, Embryological developmental (EDev) Network, constructed from the largest connected component of the developmental time-specific proteins. The size of each node is proportional to its degree, while the color represents the community it belongs to. An edge is built whether a physical interaction was reported in the human interactome.

Encouraged by this result, we collected all time-specific proteins before birth, obtaining a list of 5,495 developmental-specific proteins. Their expression in the original dataset was significantly higher during development compared to other time periods (Fig. 3B, *p-value*: 5e-270, two-tailed T-test), suggesting their transcriptomic importance. Furthermore, their connectivity in the interactome was higher than expected by chance (*z-score*: 9.5, Fig. 3C), suggesting a highly interconnected involvement of these proteins during human development. This network was enriched for essential biological mechanisms such as DNA transcription and duplication (Fig. 3D; Supplementary Table 6). Due to the highly interconnected nature of this protein network, we first defined and then partitioned the embryological developmental (EDev) network (composed of 5,349 nodes and 58,936 edges) in 11 distinct communities (Fig. 3E), corresponding to distinct biological functions (Supplementary Fig. 12; Supplementary Table 7). For example, community 5 is enriched with proteins specialized in oxidative phosphorylation, while proteins in community 7 are responsible for cell cycle regulation. Thus, the EDev network and its sub-clusters provide a detailed map of molecular activities crucial for early human development.

### Understanding the relationship between aging and the Developmental Origins of Health and Diseases (DOHaD)

To elucidate the involvement of the EDev network in disease etiology and pathogenesis we collected proteins associated with several embryological conditions, spanning congenital defects to Mendelian rare diseases and pediatric cancers ^15^ (Supplementary Table 8A). Our analysis revealed that 12 out of 34 pathological conditions are statistically over-represented in the EDev network and its communities (Fig. 4A; Supplementary Table 8B; *FDR*<0.05, Fisher’s exact test and Benjamini-Hochberg correction). Moreover, 26 of all conditions showed higher connectivity in this network than randomly expected (*z-score*>2, Fig. 4B; Supplementary Table 8B), suggesting the existence of coordinated molecular mechanisms, termed disease modules, underlying the phenotypic manifestation of these conditions and pointing to an incomplete understanding of their molecular origins. Interestingly, specific communities within the EDev network appear to be involved only in certain diseases. For instance, most common pediatric cancers are particularly enriched in community 8 (*FDR*=0.01, *z-score*=4), primarily associated with Ras signaling, the PI3K pathway, and cancer pathways. In contrast, several rare diseases and rare cancer syndromes tend to be enriched and well-connected in community 4, which is mainly involved in DNA damage and DNA repair.

**Figure 4:**
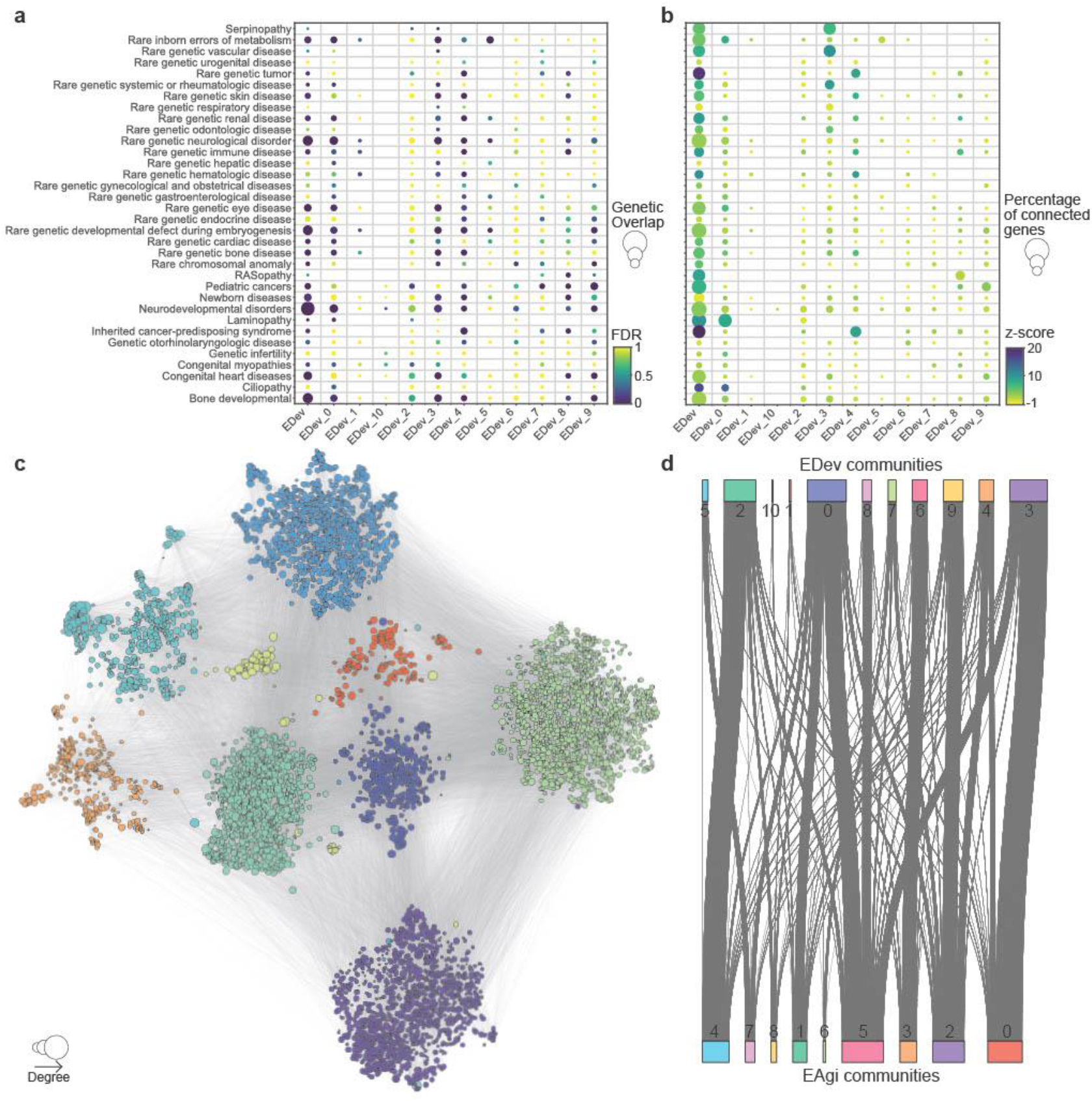
The EDev network captures the molecular changes occurring in embryological disorders. **A**, Heatmap showing the enrichment of several developmental diseases in the EDev network and its communities. The size of each circle is proportional to the number of shared proteins. **B**, Heatmap showing the connectivity z-score of the proteins associated with several developmental diseases in the EDev and its communities. The size of each circle is proportional to the percentage of connected proteins. **C**, The Environmental Aging (EAgi) Network. Each node represents a protein, its size is proportional to its degree in the network, and its color depends on the community it belongs to. An edge is drawn whether a physical interaction between two proteins was reported in the human interactome. **D**, Alluvial plot showing the rearrangement of the proteins from the communities of the EDev network to the communities of the EAgi network.

In a parallel analysis, we defined an environmental aging (EAgi) network by collecting 5,920 time-specific proteins occurring in the networks referring to postnatal time points. This network comprises 5,848 nodes and 68,929 edges (Fig. 4C), forming a well-connected neighborhood within the global interactome (*z-score*: 10, Supplementary Fig. 13A). Surprisingly, the EAgi network shows a high similarity with the EDev network, sharing 4,612 proteins (*p-value* <1e-278, hypergeometric test; Supplementary Fig. 13B), suggesting the involvement of the same proteins during environmental aging and embryonic development.

Next, we identified 9 communities in this EAgi network. Interestingly, only one community shared more than 50% similarity with any communities of the EDev network. Instead, most proteins were distributed differently in the EAgi and EDev communities (Fig. 4D), indicative of distinct molecular reprogramming between embryonic development and environmental aging. Supporting this hypothesis, we observed a broad range of molecular activities associated with each community in the EAgi network (Supplementary Fig. 14; Supplementary Table 9). Some specific communities were enriched in known age-related mechanisms, such as the histone modifications and epigenetic changes in community 4 ^16^, and the protein degradation/ubiquitination machinery in community 8 ^17^. Moreover, shared molecular functions between development and aging were identified as well. For instance, mitochondrial activity, which is crucial to provide energy to the cell, is important in both contexts, with its impairment identified in age-related processes ^18^. Ultimately, we conclude that the main difference between embryological development and environmental aging lies not in the proteins themselves, but in their coordinated activities over time. Indeed, we observed that the number of proteins involved in each community over time was statistically significantly different between pre- and post-conception for 9 out of the 11 communities in EDev and 5 out of the 9 communities in EAgi (*FDR*<0.05, two-sided independent T-test with Benjamin-Hochberg correction) (Supplementary Fig. 15A, B), with the most striking being EDev_9 (*FDR*=3e-8) and EAgi_0 (*FDR*=2e-07), representing a decreased activity of transcription events and an increased involvement of the lysosomal activity after birth, respectively.

Finally, we investigated whether these coordinated molecular programs occurred uniformly across all tissues or displayed specific organ affinities. To do this, we calculated the protein overrepresentation and the significance of connectivity in the context of the previously identified tissue-specific proteins. Overall, these communities were well represented in all tissues for both EDev and EAgi (Supplementary Fig. 16A; Supplementary Table 10A), though their connectivity patterns differed significantly (Supplementary Fig. 16B; Supplementary Table 10A). For example, the heart exhibited a distinct affinity for the EAgi community 8 (*FDR*=2e-12, *z-score*: 2.4), responsible for protein ubiquitination, which has been involved in cardiovascular diseases ^19,20^, and aging ^21^. On the other hand, communities governing basic mechanisms such as MAPK, and PI3K-Akt signaling (EAgi community 5) were statistically significantly shared across multiple tissues (brain, cerebellum, kidney, ovary, and testis). Furthermore, we observed a high correlation of the connectivity patterns across tissues both in embryonic development (Supplementary Fig. 16C; Supplementary Table 10B) and environmental aging (Supplementary Fig. 16D; Supplementary Table 10C). An exception was the liver, which showed a unique pattern of connectivity during embryonic development, consistently with the broad spectrum of biological activities that this organ executes during the fetal period (ranging from hematopoiesis to metabolism) ^22^ .

### Elucidating the impact of interventional drugs in aging

To systematically elucidate the underlying molecular mechanisms underpinning aging in our networks, we have compiled a list of 502 expert-curated proteins associated with human aging from ^23^, and grouped into 10 distinct classes (Supplementary Table 11A) based on their molecular activity. As previously performed, we tested for their protein overrepresentation (Fig. 5A; Supplementary Table 11B) and their network connectivity in the EDev and the EAgi networks (Fig. 5B; Supplementary Table 11B). Consistent with our previous findings, we observed some shared molecular mechanisms enriched in both networks (such as mitochondrial dysfunction). However, by integrating connectivity information, we were able to identify that EAgi community 5 is particularly relevant to aging. This community involves pathways such as ‘cellular senescence’ (*FDR*: 3e-05, *z-score*: 15), ‘NF-kB pathway’ (*FDR*: 5e-06, *z-score*: 18), ‘senescence-associated secretory phenotype’ (*FDR*: 4e-05, *z-score*: 4), and ‘others’ (*FDR*: 6e-08, *z-score*: 4.5) (Supplementary Table 11B). Interestingly, EAgi community 5 exhibited the highest association with proteins whose protein variants have been observed to correlate with longevity in humans ^24^ (Supplementary Fig. 17A; *p-value*: 0.003, Fisher’s exact test). Encouraged by these results, we wondered whether we could identify the mechanisms of action of anti-aging drugs that have been tested so far. To achieve this goal, we assembled a collection of 500 drugs that have shown the capability to reverse aging in animal models ^25^ and we predicted their human targets using the Comparative Toxicogenomic Database ^26^ (CTD) (Supplementary Table 12A). By examining their protein overrepresentation and connectivity patterns, we predicted which communities of the EDev and the EAgi would be targeted, finding that the highest number of candidate drugs target the EAgi community 5 (Fig. 5C; Supplementary Table 12B, C). Given that many of these drugs have multiple targets, influencing diverse aging mechanisms, we postulated that the most efficacious anti-aging drugs would target a broader spectrum of EAgi communities. Our analyses showed that each drug tends to statistically significantly target at most 2 communities of the EAgi, identifying a list of the most promising 8 candidate drugs (Fig. 5D; Supplementary Table 12D). Among the top-ranked compounds, Resveratrol emerges as a potential anti-aging molecule targeting 100 proteins in the aging context. Resveratrol is a non-flavonoid polyphenol produced by various plants, including grapes and blueberries, and is known for its antioxidant, anti-inflammatory, anticarcinogenic, and anti-aging properties ^27^. These results have been confirmed also by considering a different metric, network proximity, whose null model considers the degree distribution of the two protein lists, correcting for the power-law structure of the interactome (Supplementary Table 12E). Also in this case, Resveratrol was found to be tightly associated with EAgi communities, such as community 5 (*z-score* = 16), community 2 (*z-score* = 32), and community 6 (*z-score* = 10). To further understand the drug-target interactions, we focused on the proteins in EAgi community 5 that were statistically significantly associated with both an aging-related molecular mechanism and an anti-aging drug. This resulted in a network of 61 proteins and 120 physical interactions (Fig. 5E; Supplementary Table 12F). Interestingly, we did not observe any correlation between the number of drugs that would target a particular protein and its degree in this subnetwork (Supplementary Fig. 17B) or in the general PPI (Supplementary Fig. 17C), suggesting that a network-guided drug targeting approach could be beneficial. For instance, despite having only one physical interaction in this subnetwork (66 in the global PPI), Vascular Endothelial Growth Factor A (VEGFA) was targeted by 33 anti-aging drugs. Conversely, only one drug is designed to modulate the activity of the most connected protein in this subnetwork, Bruton’s tyrosine kinase (BTK), which has 10 interactions (144 in the general PPI). BTK has been recently shown to be associated with aging and its inhibition reduced the number of senescent cells in tissues in mice ^28^. In summary, our network-driven approach provides a comprehensive framework for understanding the mechanisms through which anti-aging drugs exert their effects, highlighting potential targets and therapeutic strategies that could mitigate aging and promote longevity.

**Figure 5:**
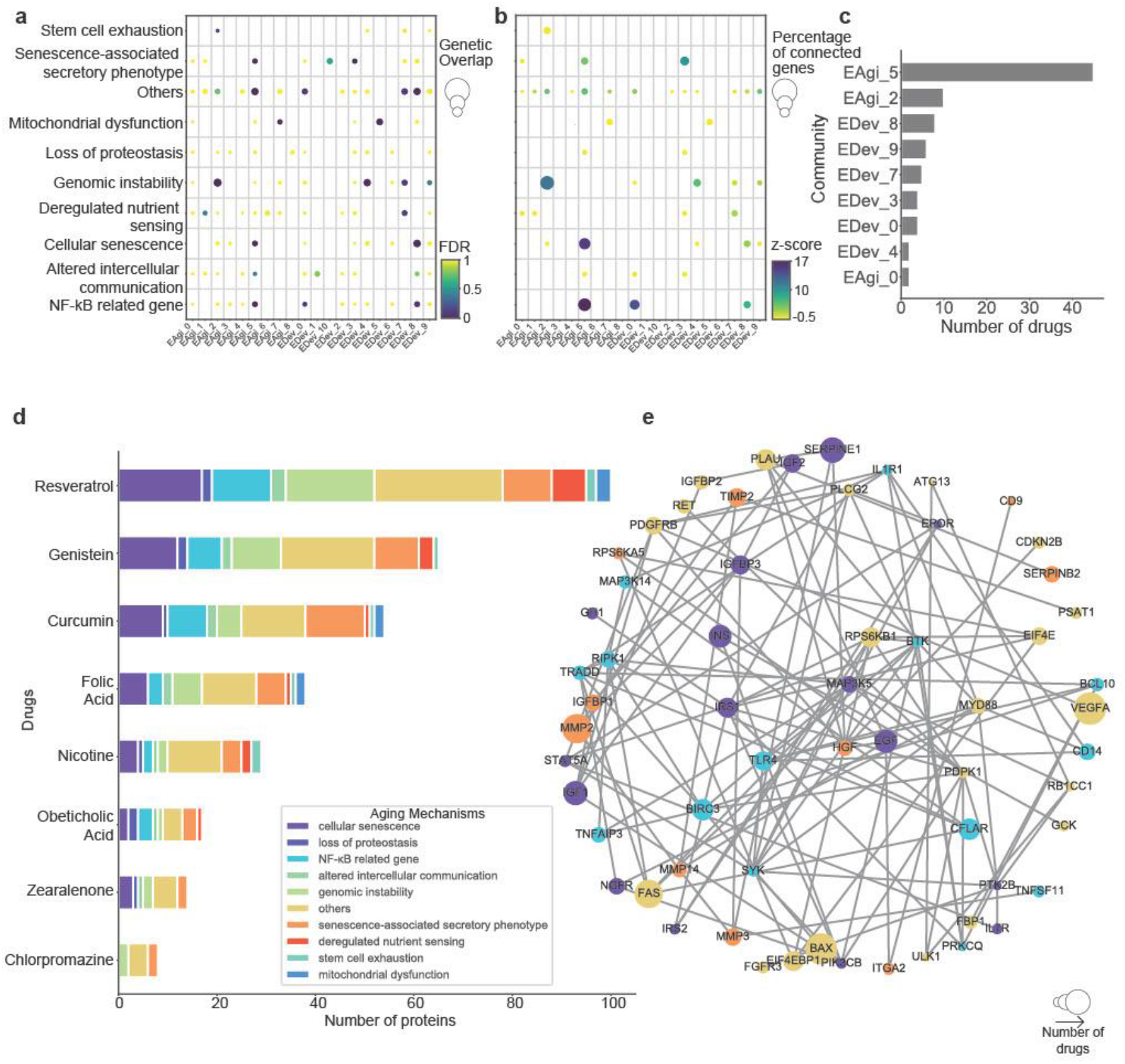
The identification of a draggable-aging subnetwork. **A**, Heatmap showing the enrichment of 10 molecular hallmarks of aging in the EDev and EAgi networks and their corresponding communities. The size of each circle is proportional to the number of shared proteins. **B**, Heatmap showing the enrichment of 10 molecular hallmarks of aging in the EDev and EAgi networks and their corresponding communities. The size of each circle is proportional to the number of connected proteins, and its color is given by the connectivity significance z-score. **C**, Barplot showing the number of anti-aging drugs targeting each of the communities in the EDev and the EAgi networks. **D**, Barplot showing the number of associated proteins for each of the 8 anti-aging drugs that target more than one single community. The color of each bar resembles the 10 annotated molecular hallmarks of aging. **E**, Anti-aging subnetwork built from the druggable proteins of the EAgi community 5 that have been associated with at least one molecular hallmark of aging. Each node represents a protein and its size is proportional to the number of anti-aging drugs that target it. The color of each node represents the associated molecular hallmark of aging. An edge is built between two nodes if there is a physical protein interaction between them.

## Discussion

Although embryonic development and aging represent distinct biological phenomena, they share parallel molecular mechanisms at their core functionality. For instance, the dysregulation of those processes characterizing embryonic development, such as cell division and proliferation, is also present in aging-related features, and often described phenotypically as cellular senescence. Similar comparisons can be held for other aging-related mechanisms, such as mitochondrial dysfunction and cellular signaling.

Despite the intuitive metaphor of this parallelism, a systematic molecular overview of these two phenomena has been lacking. In this study, we used transcriptomic data from seven distinct tissues across different life stages to construct and trace the evolution of the underlying molecular networks that regulate each phase of life. From our analysis, we identified a set of proteins expressed consistently across all tissues and time points, which we termed *common-core*. These proteins are primarily involved in housekeeping activities essential for cell survival and tend to interact with each other, forming densely connected protein complexes. Moreover, in agreement with their important biological roles, the common-core proteins tend to occupy central positions in the human interactome, showing a higher degree than other proteins and participating in a broad range of biological functions.

While proteins with time-specific or tissue-specific activity have lower degree values, they can be used to construct two distinct networks reflecting embryonic development (EDev) and environmental aging (EAgi). Despite sharing a large number of proteins, local connections (communities) within these two networks demonstrated significant differences, suggesting a rearrangement of physical interactions occurring between human development and aging. By making use of these two networks, we identified pathogenetic mechanisms during embryonic development and molecular alterations characterizing human aging, which can provide potential targets for candidate anti-aging drugs. Notably, we observed that one of the proteins with the highest number of designed anti-aging compounds, VEGFA, plays a relatively small role in the aging molecular subnetwork. In contrast, more central proteins such as BTK have a relatively low number of targeted drugs. BTK is a tyrosine kinase playing a crucial role in B cell development and the immune system response ^29^. In the context of aging, its inhibition has shown to reduce the number of senescent cells, prolonging lifespan in progeroid mice ^28^. Among the candidate anti-aging compounds, Resveratrol emerged prominently, targeting 100 proteins involved in aging, aligning with its established antioxidant, anti-inflammatory, anticarcinogenic, and anti-aging properties. Genistein and curcumin ranked as the second and third top compounds, both having anti-oxidant and beneficial properties applied in the context of aging ^30,31^. Interestingly, all top 3 predicted compounds are natural products found in food, reinforcing recent findings emphasizing the crucial role of diet in maintaining a healthy lifestyle and promoting healthy aging ^32^. Conversely, our analyses revealed some chemical compounds associated with EDev communities, such as Simvastatin (community 0 and 9), Ethanol (community 3), and Floxuridine (community 4) which are indeed not recommended molecules during pregnancy due to their known potential for causing fetal aberrations ^33–35^. A commonly used drug for terminating pregnancy, Mifepristone, was predicted to be enriched in community 3 of the EDev ^36^.

The main limitation of this study lies in the small number of samples, which limits the exploration of individual differences and diminishes the statistical power. To compensate for this, we have based our protein selection criteria on GTEx data, increasing the number of samples used for selecting highly expressed proteins. Additionally, the imbalance in the number of pre-conception and post-conception samples could affect the performance of our analysis. Another limitation is that our study is based solely on transcriptomic data, and most of the network-level analyses were conducted within the context of protein-protein interaction networks, ignoring potential post-transcriptional regulatory mechanisms. We have focused mainly on PPIs, but other types of biological networks could be constructed such as gene regulatory networks (GRNs) and co-expression networks, providing additional and complementary biological contextualization. Despite these limitations, this study represents the first systematic and unbiased molecular comparison between development and aging, offering valuable insights and laying the groundwork for further investigation into analyzing degenerative diseases and aging.

## Materials & Methods

### Data sample collection and processing

Bulk RNA-seq samples of 7 distinct tissues (brain, cerebellum, heart, kidney, liver, ovary, and testis) were collected from ^8^ following their temporal evolution for 21 time points (13 samples in gestational times and 7 after birth). Specifically, the prenatal development was followed from 4 weeks-post conception (wpc) to 19 wpc on a weekly basis (except for 14,15, and 17 wpc). Postnatally time points are: newborn, infants (6-9 months), toddlers (2-4 years), school-age (7-9 years), teenagers (13-19 years), young adult (25-32 years), middle adult (46-54 years), elderly (58-63 years). Sample preparation and RNA processing are extensively described in ^8^. For the aim of this study, we have used as input data the Transcript Per Kilobase Million (TPM) matrix, which normalizes for library size and it is freely available in (https://www.ebi.ac.uk/biostudies/arrayexpress/studies/E-MTAB-6814).

### Global Interactome network construction

We combined three widely used interactomes, harnessing their advantages: (i) an interactome used for identifying links between diseases ^6^, (ii) a large-scale dataset from the HIPPIE (21) database (v2.2) ^37^, (iii) and a systematic high-throughput interactome that was previously shown to be able to infer tissue-specific networks elucidating underlying molecular mechanisms of tissue-specific phenotypes ^7^. The joint interactome is larger and denser than the individual ones but follows the same network characteristics as previously published interactomes, particularly concerning the close relationship between network distance and biological function. The aggregate network consists of 18,853 nodes and 483,037 edges and we have described it in detail in ^9^.

### Tissue/Time-specific interactome network construction

To increase the number of available samples for each tissue, we have used the Genotype-Tissue Expression (GTEx) data ^38^, which provides genome-scale expression profiles across 53 human tissues of different ages. For each of the seven tissues in Cardoso-Moreira, we have downloaded the equivalent TPMs GTEx expression matrix, grouping tissues based as described in Supplementary Table 13 (i.e. for Cerebellum both “Brain Cerebellum” and “Brain Cerebellar Hemisphere” have been included). For each of the seven organs, transcripts with the top 20* percentile median protein expression within the corresponding tissue samples would be considered as GTEx-tissue-specific proteins. Next, the median expression of the GTEx-tissue-specific proteins would be calculated on the corresponding tissue matrix from Cardoso-Moreira at each time point. We have collected all proteins with a higher expression than this cut-off and used them as seed proteins for a network propagation algorithm (random walk with restart) in the global interactome ^39^, with a restarting probability, *r* = 0.9, ensuring that the propagation remained close to the original seed protein set. The output of this procedure returned a list of all proteins in the global interactome ranked by decreasing visiting probability, representing their distance to the seed protein set. Respecting this rank, we have added each protein to the list of seed proteins until all seed proteins are connected, obtaining 119 tissue/time-specific interactomes.

### Network similarity comparison and protein-groups definition

The 119 tissue/time-specific networks were compared by calculating the Jaccard Index (JI) of the edge lists for each network pair (*EA, EB*).

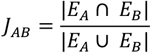

The visualization was performed by using the Python package seaborn ^40^. The 2D map was calculated by applying the SMACOF algorithm for metric multidimensional scaling in the scikit-learn Python package ^41^, and by choosing the dissimilarity measure as d_A,B_=1-*J*_*AB*_, for each pair of networks A,B.

The protein-groups definition was performed by comparing the nodes of each tissue/time-specific interactome, resulting in:

- *Common-core proteins*: nodes that are present in all 119 interactomes
- *Time-specific proteins*: nodes that are present in all interactomes of the same time-point, but excluding the common-core proteins
- *Tissue-specific proteins*: nodes that are present in all interactomes of the same tissue, but excluding the common-core proteins
- *Tissue/Time-unique proteins*: nodes that are present only in one specific tissue/time-specific interactome.

### Biological characterization of the different protein-groups

For five out of the seven tissues the highly expressed proteins, highly frequent proteins, and tissue-signature proteins were selected from an external validation set, which consisted of the single-cell RNAseq expression for brain, cerebellum, heart, liver, and kidney during development (https://descartes.brotmanbaty.org/bbi/human-protein-expression-during-development/) ^14^. For each tissue, the correlation with the mean protein expression extracted from ^8^ across all samples was calculated by a spearman correlation using the Python package Scipy. The protein name conversion from ENSEMBL IDs to protein Symbol was performed by using the Python package mygene ^42^. The highly expressed proteins were selected by using a cutoff of TPM=100 for all tissues, the most frequent proteins were calculated by taking those proteins that are expressed in at least 50% of the cells in the considered tissue, the tissue-signature proteins consisted of the differentially expressed proteins of the most common cell type in each tissue, as described in ^14^. Specifically choosing: granule neurons in cerebellum, excitatory neurons in brain, cardiomyocytes in heart, metanephric cells in kidney, and hepatoblasts in liver. All statistical comparisons with the different protein groups were performed by using both the hypergeometric test and the multinomial test. The common-core proteins were tested for their enrichment of 2176 housekeeping proteins curated from ^43^. The statistical analysis was performed using a hypergeometric test using the Python package Scipy ^44^. The list of protein complexes was downloaded from (http://humap2.proteincomplexes.org/) ^45^, which consisted of a comprehensive list of predicted protein complexes in human cells from mass spectrometry experiments. The significance of enriched protein complexes belonging to the common-core proteins was calculated by checking against a random distribution of the number of protein complexes in 1,000 protein sets of the same size as the common-core. We have curated a list of 3020 essential proteins agglomerating: 283 proteins reported to be essential in multiple cultured cell lines based on short hairpin RNA (shRNA) screening ^46^, 683 proteins reported to be essential in multiple cultured cell lines based on CRISPR/Cas screening ^47^, and 2,454 proteins which homozygous knockout in mice results in pre-, peri- or post-natal lethality ^48–50^. The comparison with all protein groups was carried out with both a hypergeometric test and a multinomial test, comparing the quantified number of overlapping proteins against the theoretical (random) expectation. To evaluate the impact of these groups of proteins on disease phenotype, we have collected a list of proteins for: Autosomal Dominant (AD), Autosomal Recessive (AR), X-linked Dominant (XD), X-linked Recessive (XR). 697 AD proteins, and 1139 AR proteins were curated from the Online Mendelian Inheritance in Man (OMIM) in ^51,52^, 31 XD proteins and 98 XR proteins in ^51^. The therapeutic component of the different regions of the PPI was performed by curating a list of FDA-approved drugs from the latest version of DrugBank (March 2023) ^53^. From our querying, we obtained a list of 904 FDA-approved drugs, which would target 988 proteins in the PPI. All statistical comparisons with the different protein groups were performed by using both the hypergeometric test and the multinomial test.

### Biological enrichment analysis

Biological enrichment was performed for all protein sets for the three main branches of the protein ontology (GO) ^54^: biological processes (BP), molecular functions (MF), and cellular components (CC), and for KEGG pathway ^55^ by using GSEAPY ^56^.

### Machine learning classifier

The classification of each time-/tissue-specific network into pre-conceptional or post-conceptional period was performed by using the time-specific proteins as predictors with 3 distinct algorithms: support vector classifier (SVC), logistic regression (LR), and random forest (RF), implemented with the Python package scikit-learn. A “leave-one-out” cross-validation approach was used.

### Network analysis and visualization

The Python package NetworkX ^57^ was used to compute basic network-based features of the 119 tissue/time-specific interactomes, such as: degree, closeness, and betweenness. Degree is defined as the number of connections (direct neighbors) that each node has on a network. The largest connected component (lcc) of a group of nodes on a network is defined as the largest subnetwork formed by the nodes belonging to the chosen group and their direct interactions. The size of the lcc of a random subset of N nodes is expected to follow a normal distribution, given that it is larger than the percolation threshold. We can therefore empirically estimate the significance of a given connected component through its z-score and corresponding empirical p-value determined from 1,000 randomly selected protein sets of the same size.

The density of the global interactome was calculated as follows:

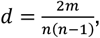

where *m* represents the number of edges and *n* the number of nodes of a network.

The network proximity analysis was performed through the python package NetMedPy ^58^, and it was used to estimate the network distance (e.g. network proximity) between the anti-aging candidate drug targets and the proteins belonging to each EDev and EAgi community.

The visualization of the Global Interactome in Fig. 1C was carried out with uniform manifold approximation and projection ^59,60^ dimensionality reduction by considering as features of the feature matrix the combination of topological network features (calculated from the visiting-probability matrix of a random-walker in the network) and the functional features (considered as community belonging for each node). The visualization of the druggable aging subnetwork in Fig. 5D was performed by calculating the spring layout of the network, which optimizes the space treating each node as a repulsive electromagnetic particle.

### EDev and EAgi networks construction

The set of 5,495 embryological developmental proteins was selected by taking the union of all time-specific proteins and tissue/time-unique proteins during the gestational period. Their median protein expression across developmental times was tested against other times by performing a two-independent sample T-test, and their network connectivity was tested by checking for their lcc, which is made of 5,349 nodes and 58,936 edges, and forms the EDev network. Similarly, the 5,920 *time-specific proteins* during the post-gestational time points, and their lcc was calculated, characterized by 5,848 nodes and 68,929 edges, namely the EAgi network.

### Network community identification

The community detection was performed by using the Louvain algorithm ^61^ built in the Python package NetworkX, and by choosing the resolution parameter that maximized network modularity. The Sankey plot visualization of the EDev/EAgi network community comparison of Fig. 4D was performed by using the Python packages plotly and kaleido.

### EDev & EAgi biological characterization

A curated list of developmental diseases and their associated proteins was collected from several sources, and presented in Supplementary Table 8A. Specifically, 8 distinct developmental conditions were selected from the “Developmental” category in the Rat Genome Database (RGD) ^62^, by filtering for only those annotations occurring in humans. An extensive curated list of 26 rare genetic disease groups from Orphanet Rare Disease Ontology was taken from ^15^. Finally, an extensive list of driver cancer proteins was extracted from a large study of pediatric cancers, analyzing the genome and transcriptome of 1,699 pediatric leukemias and solid tumors ^63^. Each of these groups of proteins were tested for significant overrepresentation in the EDev network and its communities by performing a hypergeometric test with Benjamini-Hochberg correction for multiple comparisons, and for network connectivity by calculating the z-score significance of the lcc. Similar analyses were performed by considering the tissue-specific proteins that were predicted from the tissue/time-specific interactome comparisons. A Pearson correlation was calculated between the connectivity z-score of the tissue-specific proteins across all tissues during aging and embryo development. A list of 10 distinct molecular mechanisms occurring in aging and their associated proteins was downloaded from (https://ngdc.cncb.ac.cn/aging/age_related_proteins) ^23^ and presented in Supplementary Table 11A. Their test for protein overrepresentation and network connectivity significance was conducted in both EDev and EAgi and their respective communities (Supplementary Table 11B). The identification of the statistically differentially represented EDev and EAgi communities was carried out by calculating the ratio of time-specific proteins for each community at each time point and grouping them in pre- and post-conceptional periods. These two distributions were compared with a two-independent sample T-test for each community, and each p-value was corrected for multiple comparison by using the Benjamini-Hochberg method.

### Anti-aging drugs effect prediction

A list of anti-aging drugs was downloaded from (https://genomics.senescence.info/drugs/) ^25^, a manual-curated database containing an extensive compilation of drugs, compounds and supplements with anti-aging properties tested in model organisms. The human genetic signature was predicted for 326 of these drugs by using the Comparative Toxicogenomics Database ^26^, and it is presented in Supplementary Table 12A. Their effect on the EDev and the EAgi was tested by performing an overrepresentation test (hypergeometric test with Benjamini-Hochberg correction), and lcc z-score calculation (Supplementary Table 12B). The longevity-associated proteins were downloaded from (https://genomics.senescence.info/longevity/) ^24^, consisting of a database containing human genetic variants associated with longevity.

## Code Availability

Code for reproducing the analyses in this article can be found at GitHub: https://github.com/superlsd/TemTisNet.

## Authors contributions

Conceptualization: J.M., S.D.L. Methodology: J.M., A.R., and S.D.L. Investigation: S.D.L. Visualization: S.D.L. Supervision: J.M. and A.R. Writing: S.D.L., A.R., and J.M.

## References

1. Silva-García, C. G. Devo-aging: Intersections between development and aging. GeroScience 45, 2145–2159 (2023).

2. López-Otín, C., Blasco, M. A., Partridge, L., Serrano, M. & Kroemer, G. Hallmarks of aging: An expanding universe. Cell (2023) doi:10.1016/j.cell.2022.11.001.

3. Xu, Y. et al. A single-cell transcriptome atlas profiles early organogenesis in human embryos. Nat. Cell Biol. 25, 604–615 (2023).

4. Palmer, D., Fabris, F., Doherty, A., Freitas, A. A. & de Magalhães, J. P. Ageing transcriptome meta-analysis reveals similarities and differences between key mammalian tissues. Aging (Albany NY) 13, 3313–3341 (2021).

5. Rolland, T. et al. A proteome-scale map of the human interactome network. Cell 159, 1212–1226 (2014).

6. Menche, J. et al. Disease networks. Uncovering disease-disease relationships through the incomplete interactome. Science 347, 1257601 (2015).

7. Luck, K. et al. A reference map of the human binary protein interactome. Nature 580, 402–408 (2020).

8. Cardoso-Moreira, M. et al. Gene expression across mammalian organ development. Nature 571, 505–509 (2019).

9. Guthrie, J. et al. AutoCore: A network-based definition of the core module of human autoimmunity and autoinflammation. Sci Adv 9, eadg6375 (2023).

10. Frenk, S. & Houseley, J. Gene expression hallmarks of cellular ageing. Biogerontology 19, 547–566 (2018).

11. Jeong, H., Mason, S. P., Barabási, A. L. & Oltvai, Z. N. Lethality and centrality in protein networks. Nature 411, 41–42 (2001).

12. Jonsson, P. F. & Bates, P. A. Global topological features of cancer proteins in the human interactome. Bioinformatics 22, 2291–2297 (2006).

13. Carbone, L. et al. Non-invasive prenatal testing: Current perspectives and future challenges. Genes (Basel) 12, 15 (2020).

14. Cao, J. et al. A human cell atlas of fetal gene expression. Science 370, (2020).

15. Buphamalai, P., Kokotovic, T., Nagy, V. & Menche, J. Network analysis reveals rare disease signatures across multiple levels of biological organization. Nat. Commun. 12, 6306 (2021).

16. Wang, K. et al. Epigenetic regulation of aging: implications for interventions of aging and diseases. Signal Transduct Target Ther 7, 374 (2022).

17. Koyuncu, S. et al. Rewiring of the ubiquitinated proteome determines ageing in C. elegans. Nature 596, 285–290 (2021).

18. Picca, A., Faitg, J., Auwerx, J., Ferrucci, L. & D’Amico, D. Mitophagy in human health, ageing and disease. Nat Metab 5, 2047–2061 (2023).

19. Drews, O. & Taegtmeyer, H. Targeting the ubiquitin-proteasome system in heart disease: the basis for new therapeutic strategies. Antioxid. Redox Signal. 21, 2322–2343 (2014).

20. Zolk, O., Schenke, C. & Sarikas, A. The ubiquitin-proteasome system: focus on the heart. Cardiovasc. Res. 70, 410–421 (2006).

21. Sosnowska, D. et al. A heart that beats for 500 years: age-related changes in cardiac proteasome activity, oxidative protein damage and expression of heat shock proteins, inflammatory factors, and mitochondrial complexes in Arctica islandica, the longest-living noncolonial animal. J. Gerontol. A Biol. Sci. Med. Sci. 69, 1448–1461 (2014).

22. Duncan, S. A. Mechanisms controlling early development of the liver. Mech. Dev. 120, 19–33 (2003).

23. Aging Atlas Consortium. Aging Atlas: a multi-omics database for aging biology. Nucleic Acids Res. 49, D825–D830 (2021).

24. Budovsky, A. et al. LongevityMap: a database of human genetic variants associated with longevity. Trends Genet. 29, 559–560 (2013).

25. Barardo, D. et al. The DrugAge database of aging-related drugs. Aging Cell 16, 594–597 (2017).

26. Davis, A. P. et al. Comparative Toxicogenomics Database (CTD): update 2021. Nucleic Acids Res. 49, D1138–D1143 (2021).

27. Salehi, B. et al. Resveratrol: A Double-Edged Sword in Health Benefits. Biomedicines 6, (2018).

28. Ekpenyong-Akiba, A. E. et al. Amelioration of age-related brain function decline by Bruton’s tyrosine kinase inhibition. Aging Cell 19, e13079 (2020).

29. McDonald, C., Xanthopoulos, C. & Kostareli, E. The role of Bruton’s tyrosine kinase in the immune system and disease. Immunology 164, 722–736 (2021).

30. Mas-Bargues, C., Borrás, C. & Viña, J. Genistein, a tool for geroscience. Mech. Ageing Dev. 204, 111665 (2022).

31. Zia, A., Farkhondeh, T., Pourbagher-Shahri, A. M. & Samarghandian, S. The role of curcumin in aging and senescence: Molecular mechanisms. Biomed. Pharmacother. 134, 111119 (2021).

32. Yeung, S. S. Y., Kwan, M. & Woo, J. Healthy Diet for Healthy Aging. Nutrients 13, (2021).

33. Karalis, D. G., Hill, A. N., Clifton, S. & Wild, R. A. The risks of statin use in pregnancy: A systematic review. J. Clin. Lipidol. 10, 1081–1090 (2016).

34. Briggs, G. G., Freeman, R. K. & Yaffe, S. J. Drugs in Pregnancy and Lactation: A Reference Guide to Fetal and Neonatal Risk. (Lippincott Williams & Wilkins, 2012).

35. Sanvisens, A. et al. Alcohol Consumption during Pregnancy: Analysis of Two Direct Metabolites of Ethanol in Meconium. Int. J. Mol. Sci. 17, 417 (2016).

36. Institute of Medicine & Committee on Antiprogestins: Assessing the Science. Clinical Applications of Mifepristone (RU486) and Other Antiprogestins: Assessing the Science and Recommending a Research Agenda. (National Academies Press, 1993).

37. Alanis-Lobato, G., Andrade-Navarro, M. A. & Schaefer, M. H. HIPPIE v2.0: enhancing meaningfulness and reliability of protein-protein interaction networks. Nucleic Acids Res. 45, D408– D414 (2017).

38. GTEx Consortium. The Genotype-Tissue Expression (GTEx) project. Nat. Genet. 45, 580–585 (2013).

39. Jacquemin, T. & Jiang, R. Walking on a tissue-specific disease-protein-complex heterogeneous network for the discovery of disease-related protein complexes. Biomed Res. Int. 2013, 732650 (2013).

40. Waskom, M. seaborn: statistical data visualization. J. Open Source Softw. 6, 3021 (2021).

41. Pedregosa, F. et al. Scikit-learn: Machine Learning in Python. arXiv [cs.LG] 2825–2830 (2012).

42. Lelong, S. et al. BioThings SDK: a toolkit for building high-performance data APIs in biomedical research. Bioinformatics 38, 2077–2079 (2022).

43. Hounkpe, B. W., Chenou, F., de Lima, F. & De Paula, E. V. HRT Atlas v1.0 database: redefining human and mouse housekeeping genes and candidate reference transcripts by mining massive RNA-seq datasets. Nucleic Acids Res. 49, D947–D955 (2021).

44. Virtanen, P. et al. SciPy 1.0: fundamental algorithms for scientific computing in Python. Nat. Methods 17, 261–272 (2020).

45. Drew, K., Wallingford, J. B. & Marcotte, E. M. hu.MAP 2.0: integration of over 15,000 proteomic experiments builds a global compendium of human multiprotein assemblies. Mol. Syst. Biol. 17, e10016 (2021).

46. Hart, T., Brown, K. R., Sircoulomb, F., Rottapel, R. & Moffat, J. Measuring error rates in genomic perturbation screens: gold standards for human functional genomics. Mol. Syst. Biol. 10, 733 (2014).

47. Hart, T. et al. Evaluation and Design of Genome-Wide CRISPR/SpCas9 Knockout Screens. G3 7, 2719–2727 (2017).

48. Liu, X., Jian, X. & Boerwinkle, E. dbNSFP v2.0: a database of human non-synonymous SNVs and their functional predictions and annotations. Hum. Mutat. 34, E2393–402 (2013).

49. Georgi, B., Voight, B. F. & Bućan, M. From mouse to human: evolutionary genomics analysis of human orthologs of essential genes. PLoS Genet. 9, e1003484 (2013).

50. Blake, J. A. et al. The Mouse Genome Database (MGD): premier model organism resource for mammalian genomics and genetics. Nucleic Acids Res. 39, D842–8 (2011).

51. Berg, J. S. et al. An informatics approach to analyzing the incidentalome. Genet. Med. 15, 36–44 (2013).

52. Blekhman, R. et al. Natural selection on genes that underlie human disease susceptibility. Curr. Biol. 18, 883–889 (2008).

53. Wishart, D. S. et al. DrugBank 5.0: a major update to the DrugBank database for 2018. Nucleic Acids Res. 46, D1074–D1082 (2018).

54. Ashburner, M. et al. Gene Ontology: tool for the unification of biology. Nat. Genet. 25, 25–29 (2000).

55. Kanehisa, M. & Goto, S. KEGG: kyoto encyclopedia of genes and genomes. Nucleic Acids Res. 28, 27–30 (2000).

56. Fang, Z., Liu, X. & Peltz, G. GSEApy: a comprehensive package for performing gene set enrichment analysis in Python. Bioinformatics 39, (2023).

57. Hagberg, A., Swart, P. J. & Schult, D. A. Exploring Network Structure, Dynamics, and Function Using NetworkX. https://www.osti.gov/biblio/960616 (2008).

58. Aldana, A. et al. NetMedPy: A python package for large-scale Network Medicine screening. bioRxivorg (2024) doi:10.1101/2024.09.05.611537.

59. McInnes, L., Healy, J., Saul, N. & Großberger, L. UMAP: Uniform Manifold Approximation and Projection. Journal of Open Source Software vol. 3 861 Preprint at 10.21105/joss.00861 (2018).

60. Hütter, C. V. R., Sin, C., Müller, F. & Menche, J. Network cartographs for interpretable visualizations. Nature Computational Science 2, 84–89 (2022).

61. Blondel, V. D., Guillaume, J.-L., Lambiotte, R. & Lefebvre, E. Fast unfolding of communities in large networks. J. Stat. Mech. 2008, P10008 (2008).

62. Vedi, M. et al. 2022 updates to the Rat Genome Database: a Findable, Accessible, Interoperable, and Reusable (FAIR) resource. Genetics 224, (2023).

63. Ma, X. et al. Pan-cancer genome and transcriptome analyses of 1,699 paediatric leukaemias and solid tumours. Nature 555, 371–376 (2018).

