## Supplementary Figures for "A multilayer network approach elucidates time- and tissue-specific developmental and aging processes"

### Supplementary Figure 1

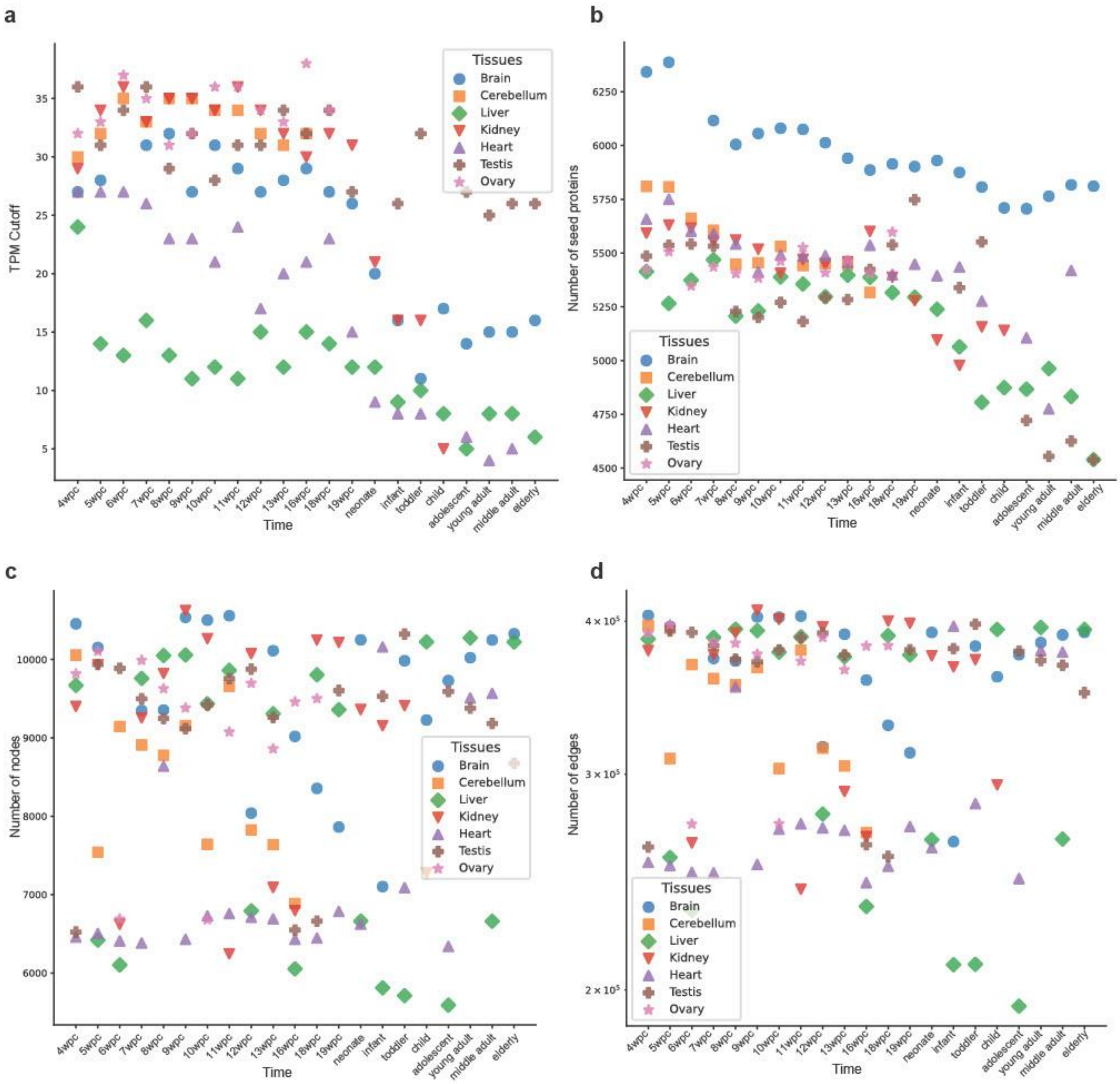


**Supplementary Figure 1**: **a**) Scatter plot showing the Transcripts per Million (TPM) cutoff for each tissue over time. **b**) Scatter plot showing the number of seed proteins for each tissue over time. **c**) Scatter plot showing the number of nodes for each time-/tissue- specific over time. **d**) Scatter plot showing the number of edges for each time-/tissue- specific over time.

### Supplementary Figure 2

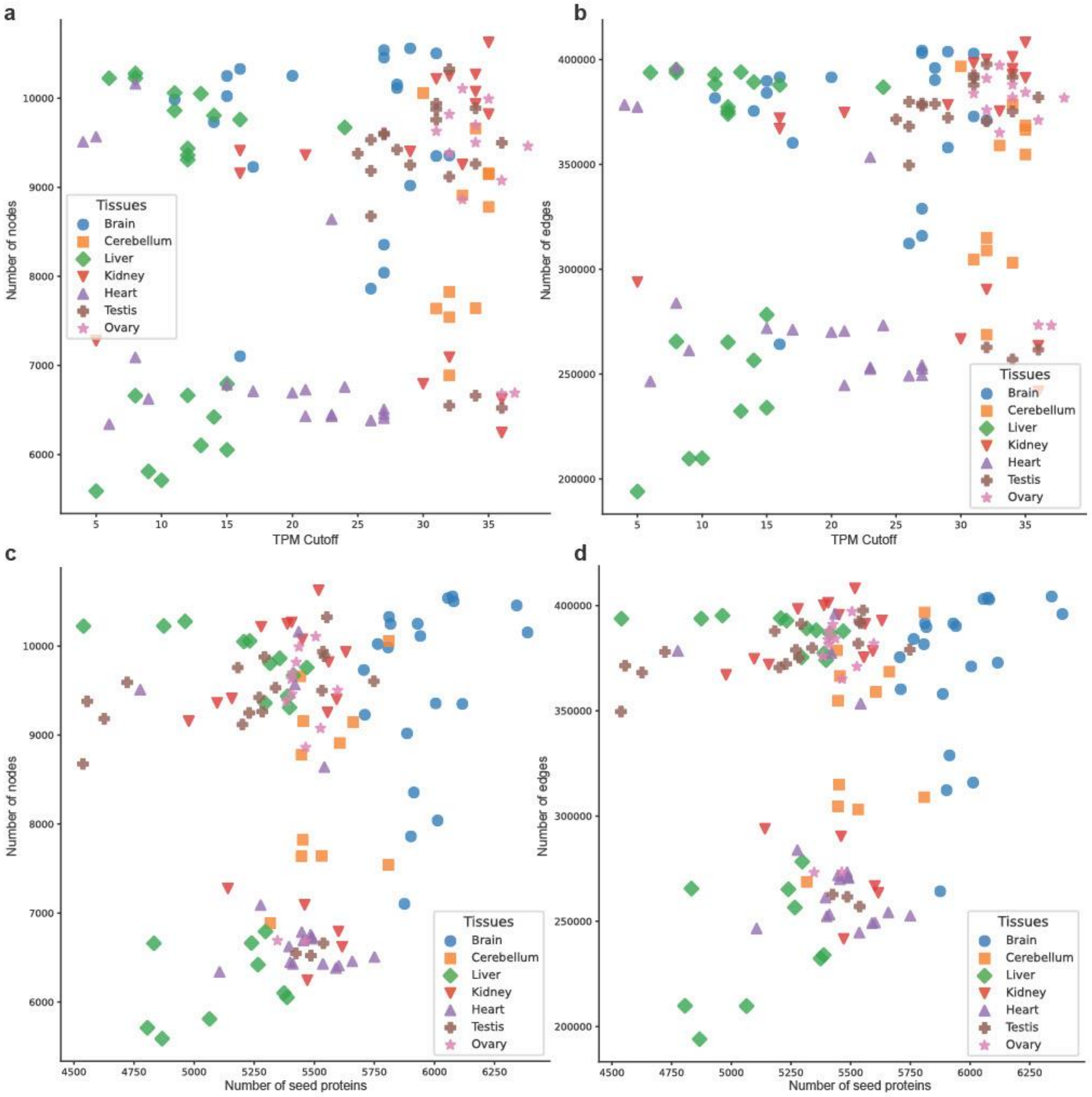


**Supplementary Figure 2**: **a**) Scatter plot showing the relationship between the number of nodes for each time-/tissue- specific network and the Transcripts per Million (TPM) cutoff. **b**) Scatter plot showing the relationship of the number of edges for each time-/tissue- specific network and the Transcripts per Million (TPM) cutoff. **c**) Scatter plot showing the relationship of the number of nodes for each time-/tissue-specific network and the number of seed proteins. **d**) Scatter plot showing the relationship between the number of edges for each time-/tissue- specific network and the number of seed proteins.

### Supplementary Figure 3

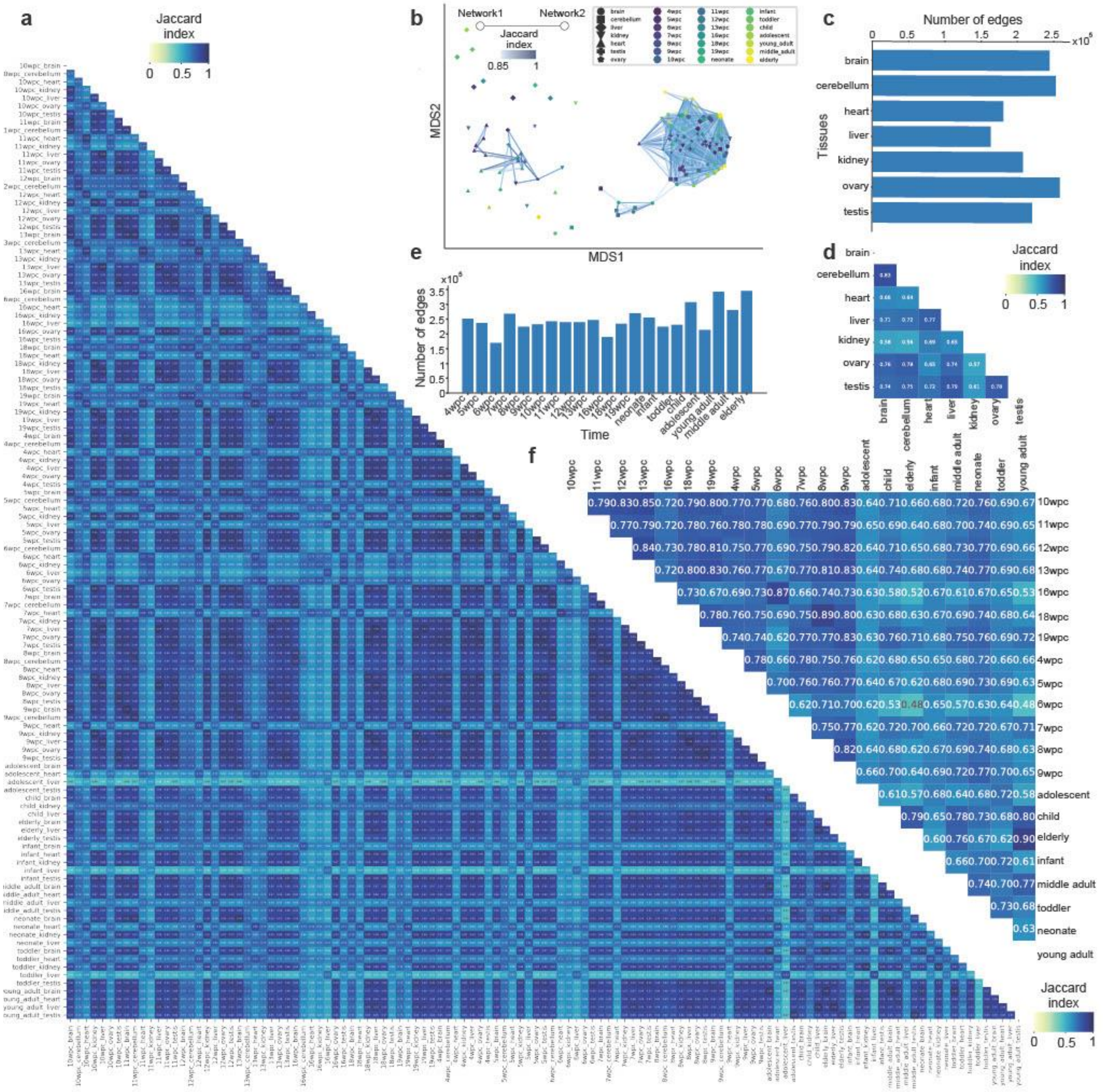


**Supplementary Figure 3**: **a**) Heatmap showing the edge similarity across all pairwise comparisons of the 119 time-/tissue- specific networks, calculated using the Jaccard Index (JI). **b**) Multidimensional scaling plot illustrating how networks derived from the same tissue or closer in time tend to cluster together. Each dot represents a tissue-/time- specific PPI network, each tissue is represented by a different shape, the gradient color follows the temporal evolution (from dark blue to light yellow). An edge is drawn between two networks, if they share at least 85% of their edges and their width is proportional to the edge overlap between the two networks. **c**) Bar plot showing the number of shared edges across all networks belonging to the same tissue. **d**) Heatmap showing tissue similarity based on edge overlap across tissue pairs, calculated using the Jaccard Index (JI). **e**)

Bar plot showing the number of shared edges across all networks belonging to the same time. **f**) Heatmap showing time similarity based on edge overlap across time pairs, calculated using Jaccard Index (JI).

### Supplementary Figure 4

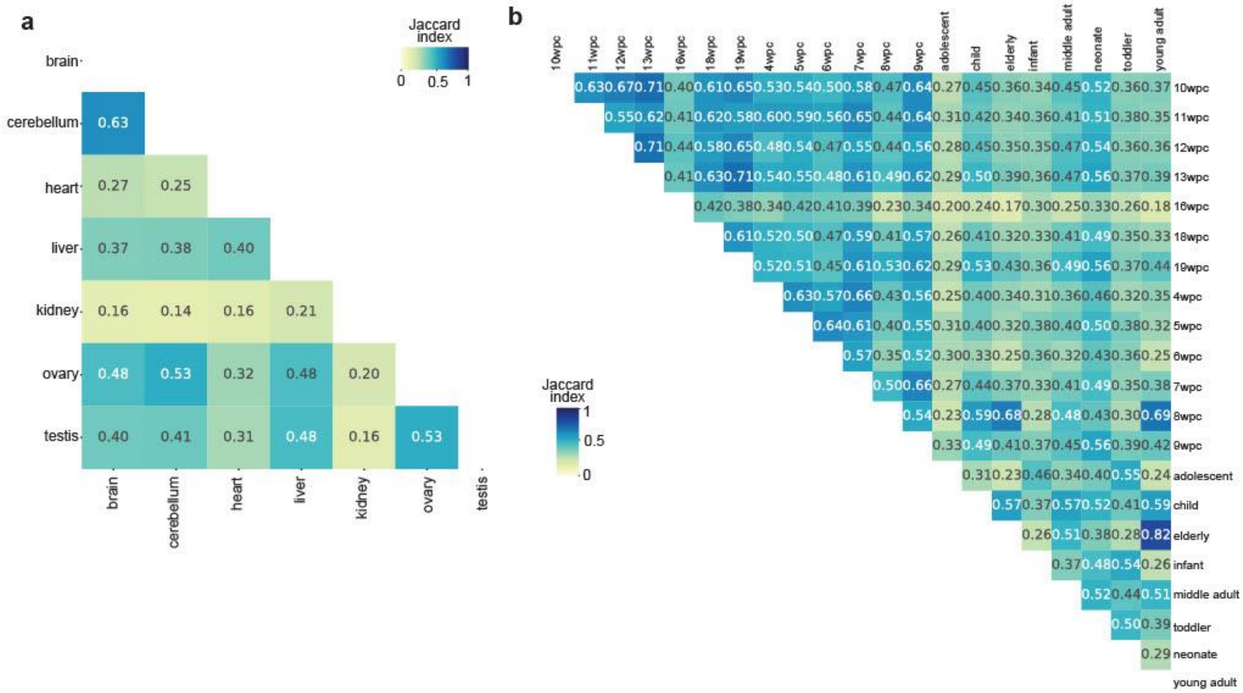


**Supplementary Figure 4**: **a**) Heatmap showing tissue similarity based on the overlap of the tissue-specific proteins, calculated using the Jaccard Index (JI). **b**) Heatmap showing time similarity based on the overlap of the time-specific proteins, calculated using the Jaccard Index (JI).

### Supplementary Figure 5

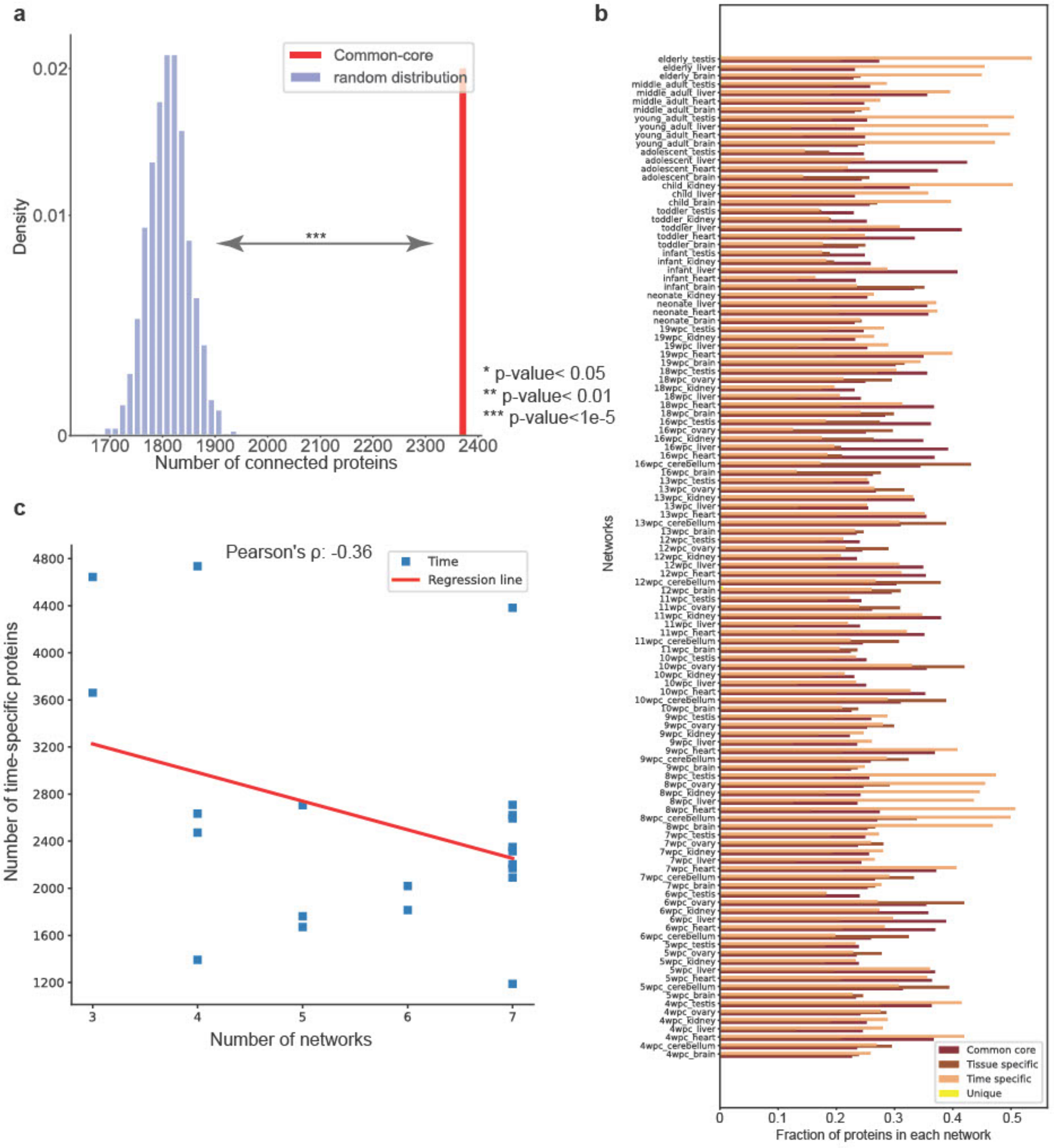


**Supplementary Figure 5**: **a**) Comparison of the connectivity in the interactome of the largest connected component of the common-core proteins against 1,000 randomized protein sets of the same size. **b**) Bar plot showing the proportion of each protein category (common-core (red), tissue-specific (brown), time-specific (orange), unique (yellow)) across all 119 time-/tissue- specific networks. **c**) Scatter plot showing the negative correlation between the number of time-specific proteins and the number of networks available for the corresponding time point.

### Supplementary Figure 6

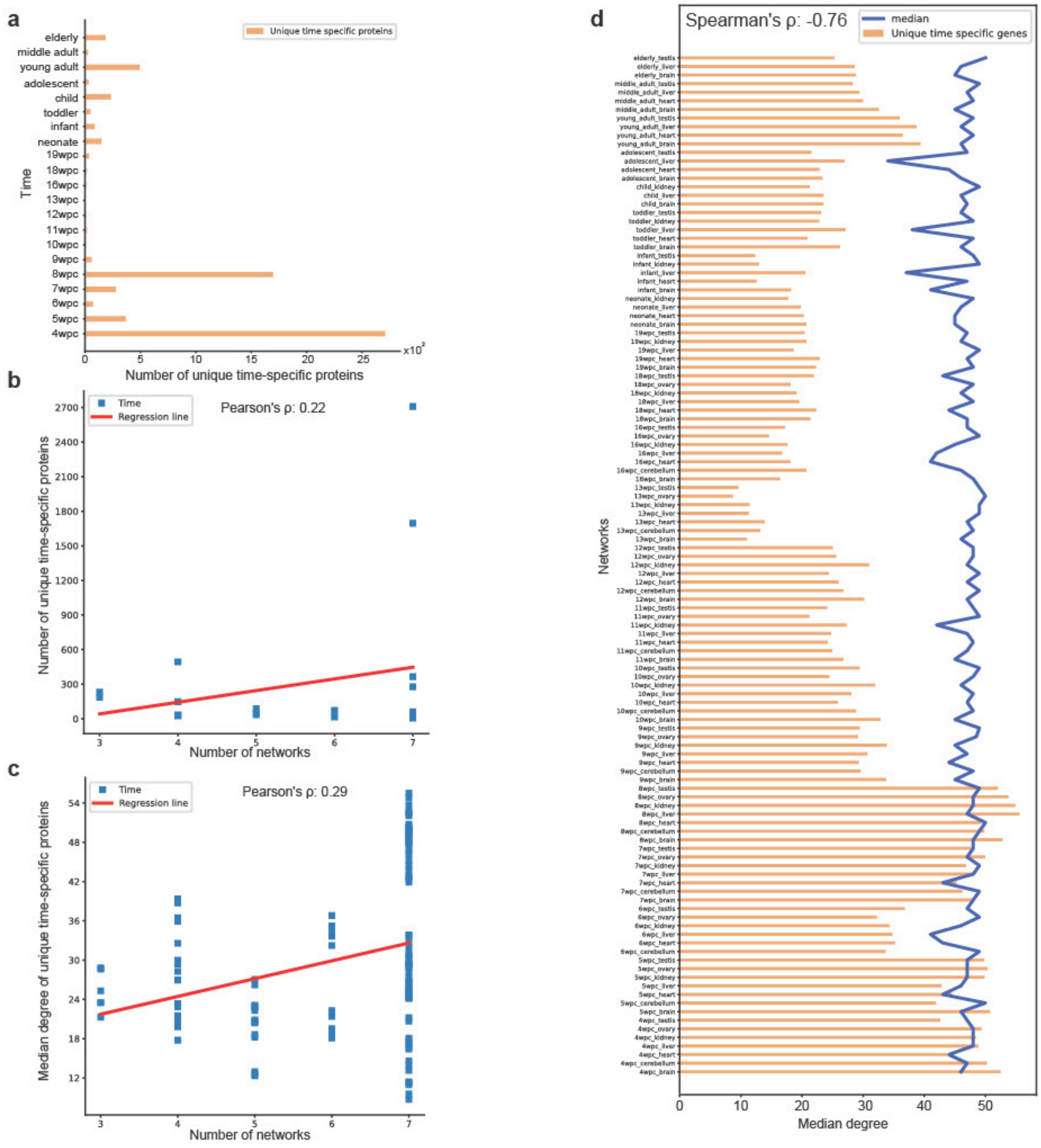


**Supplementary Figure 6**: **a**) Bar plot illustrating the number of unique time-specific proteins (time-specific proteins occurring for the first time during the lifespan) at each time point. **b**) Scatterplot showing the correlation between the number of unique time-specific proteins and the number of available networks at the corresponding time point. **c**) Scatter plot showing the correlation between the median degree in the interactome of unique time-specific proteins and the number of available networks in the corresponding time point. **d**) Bar plot showing the progression of the median degree of unique time-specific proteins (yellow bars) and the overall median degree of all proteins (blue line) over time across all 119 time-/tissue- specific networks.

### Supplementary Figure 7

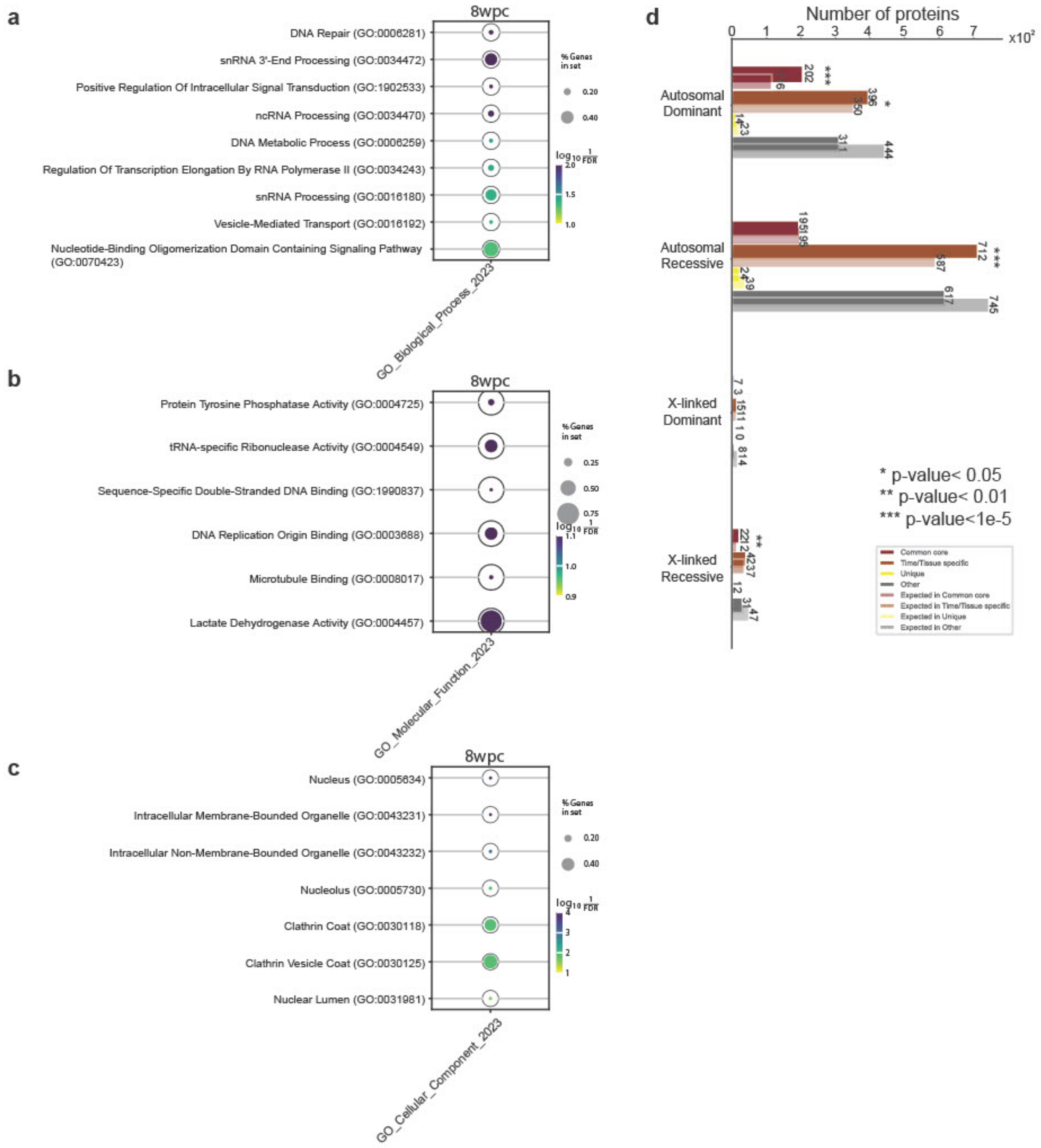


**Supplementary Figure 7**: Biological enrichment of the unique time-specific proteins corresponding at the 8th week post conception (wpc) for the three branches of the Gene Ontology: Biological Processes (BP) (**a**), Molecular Functions (MF) (**b**), and Cellular Components (CC) (**c**). **d**) Bar plot showing the number of expected and observed proteins enriched in: common-core (red), time- or tissue-specific (brown), time-/tissue unique (yellow) and other (grey) for proteins associated with mendelian diseases (autosomal dominant and recessive, and X-linked dominant and recessive). The statistical significance was calculated through Fisher's exact test.

#### Supplementary Figure 8

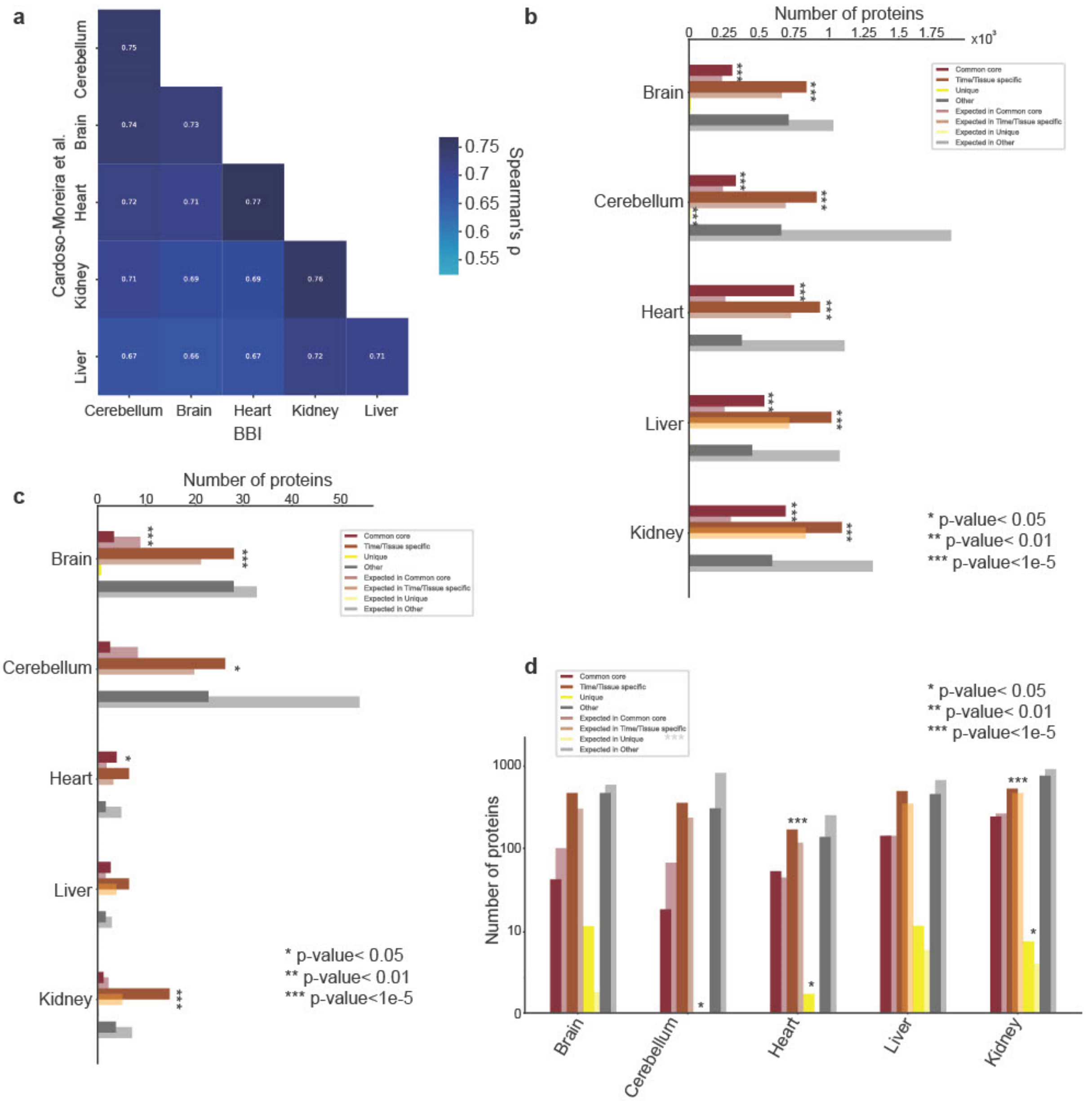


**Supplementary Figure 8**: **a**) Heatmap showing the Spearmean's rho correlation value of the median expression between the transcripts in the bulk RNA-seq experiment in Cardoso Moreira et al. and the single-cell RNA-seq in BBI. **b**) Bar plot showing the number of expected and observed highly expressed proteins for each tissue identified in BBI enriched in: common-core (red), time- or tissue-specific (brown), time-/tissue unique (yellow) and other (grey). **c**) Bar plot showing the number of expected and observed the most frequently expressed proteins for each tissue identified in BBI enriched in: common-core (red), time- or tissue-specific (brown), time-/tissue unique (yellow) and other (grey). **d**) Bar plot showing the number of expected and observed tissue-signature proteins for each tissue identified in BBI enriched in:

common-core (red), time- or tissue-specific (brown), time-/tissue unique (yellow) and other (grey). The statistical significance in b-d was calculated through Fisher's exact test.

### Supplementary Figure 9

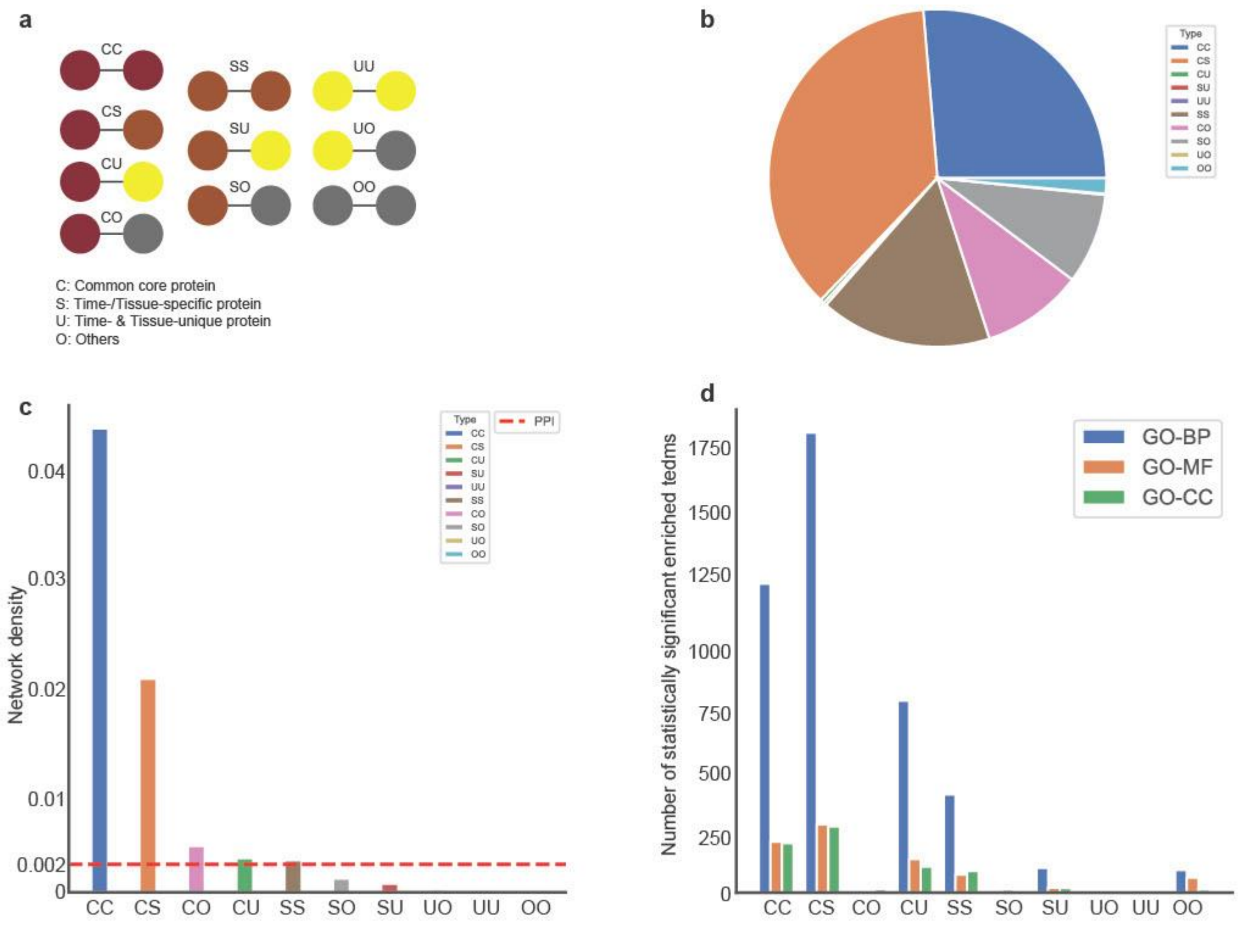


**Supplementary Figure 9**: **a**) Illustration of the 10 possible types of interactions occurring in the PPI network. **b**) Pie-chart showing the proportion of each of the 10 types of interactions in the PPI network. **c**) Barplot showing the network density of the proteins belonging to each of the 10 types of interactions. A threshold (dashed-red line) shows the network density of the PPI network. **d**) Number of statistically significant enriched terms (FDR<0.05) for the set of proteins that are part of the 10 types of interactions in the PPI. The enrichment was performed on the protein Ontology (GO) for Biological Processes (BP), Molecular Functions (MF), and Cellular Components (CC).

### Supplementary Figure 10

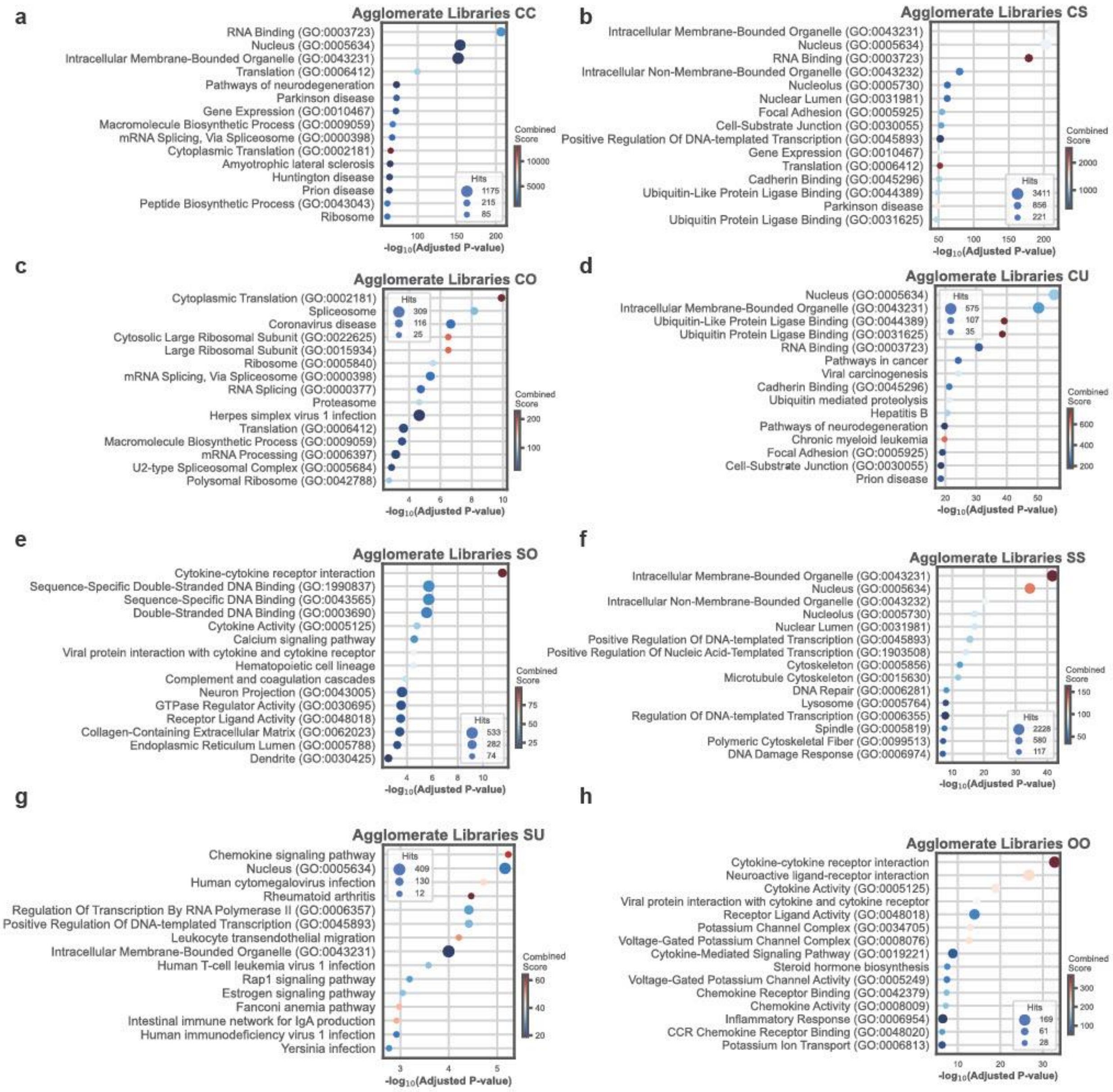


**Supplementary Figure 10**: Biological enrichment for the three branches of the Gene Ontology: Biological Processes (BP), Molecular Functions (MF), and Cellular Components, and the KEGG pathway for the genes involved in the following edges: common core – common core (CC, **a**); common core – specific (CS, **b**); common core – others (CO, **c**); common core – unique (CU, **d**); specific – others (SO, **e**); specific – specific (SS, **f**); specific – unique (SU, **g**); others – others (OO, **h**).

### Supplementary Figure 11

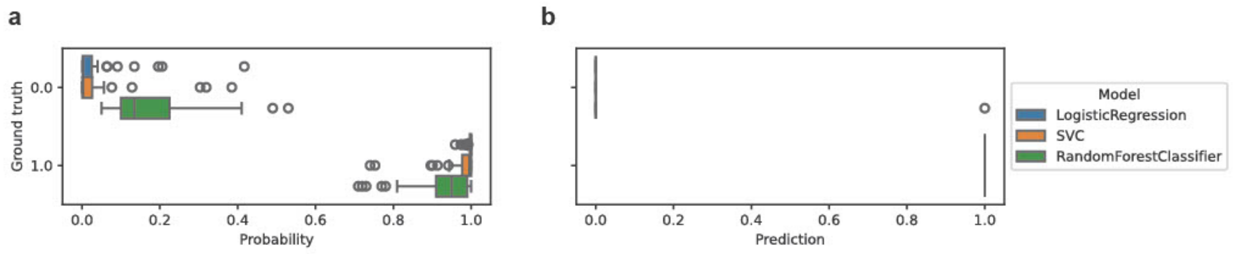


**Supplementary Figure 11**: **a**) Box plot showing the classification accuracy of predicting each network's proper time (pre-conception = 1, post-conception = 0) from the time-specific proteins by using three machine learning models: Logistic Regression (blue), Support Vector Classifier (orange), and Random Forest (green). **b**) Agglomerated prediction scores for the two classes (pre-conception = 1, post-conception = 0).

### Supplementary Figure 12

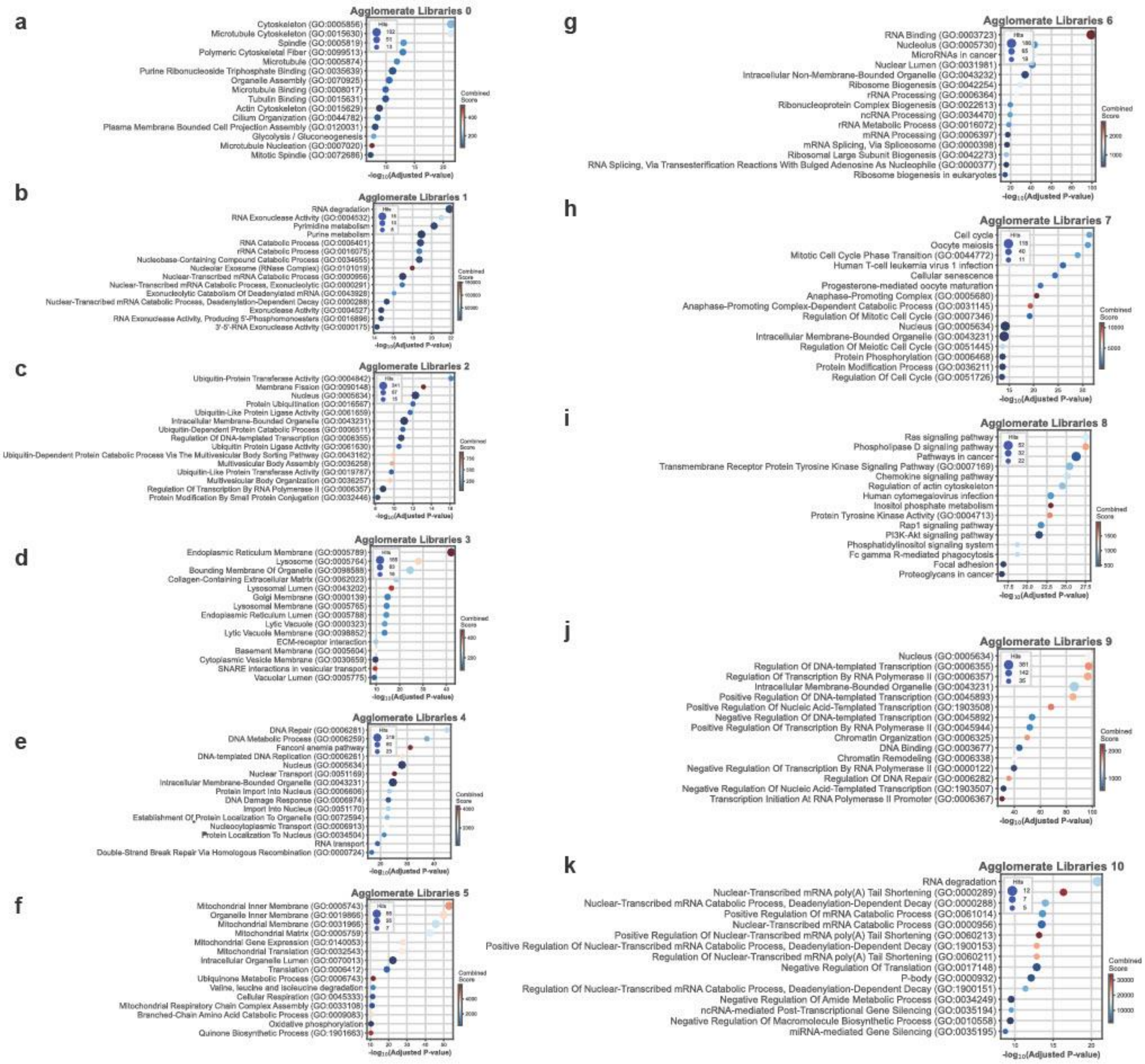


**Supplementary Figure 12**: Biological enrichment for the three branches of the Gene Ontology: Biological Processes (BP), Molecular Functions (MF), and Cellular Components, and the KEGG pathway for the genes of the 11 Embryological Developmental (EDev) network communities (**a-k**).

### Supplementary Figure 13

a

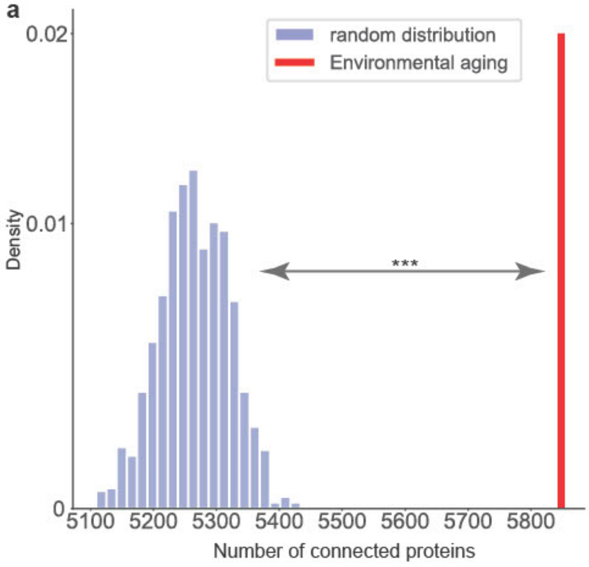


b

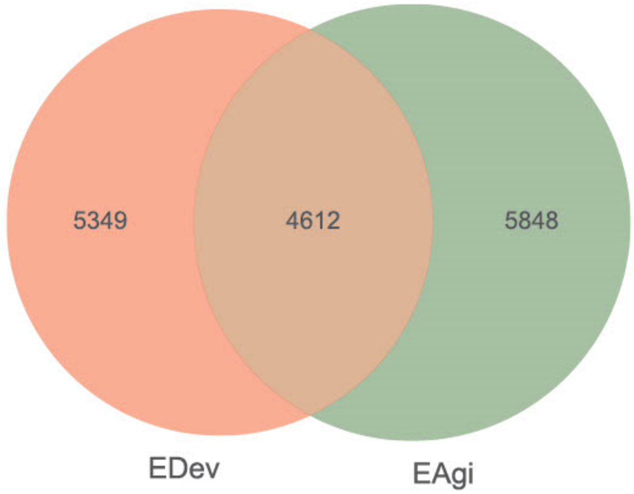


**Supplementary Figure 13**: **a**) Comparison of the connectivity in the interactome of the largest connected component of the Environmental Aging (EAgi) proteins against 1,000 randomized protein sets of the same size. **b**) Venn-Diagram showing the overlap between the Embryological Development (EDev) network proteins and the Environmental Aging (EAgi) network proteins.

### Supplementary Figure 14

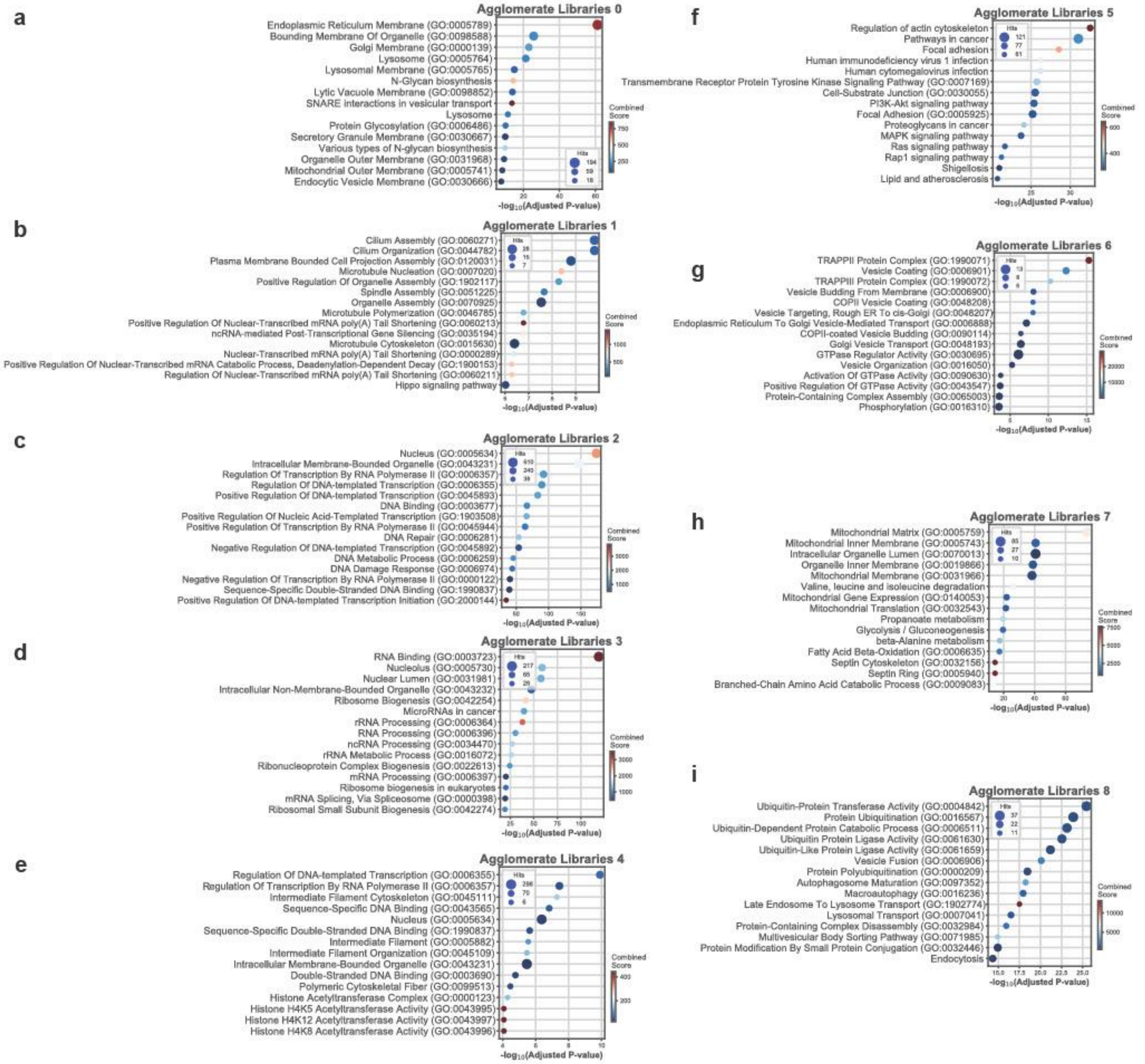


**Supplementary Figure 14**: Biological enrichment for the three branches of the Gene Ontology: Biological Processes (BP), Molecular Functions (MF), and Cellular Components, and the KEGG pathway for the genes of the 9 Environmental Aging (EAgi) network communities (**a-i**).

### Supplementary Figure 15

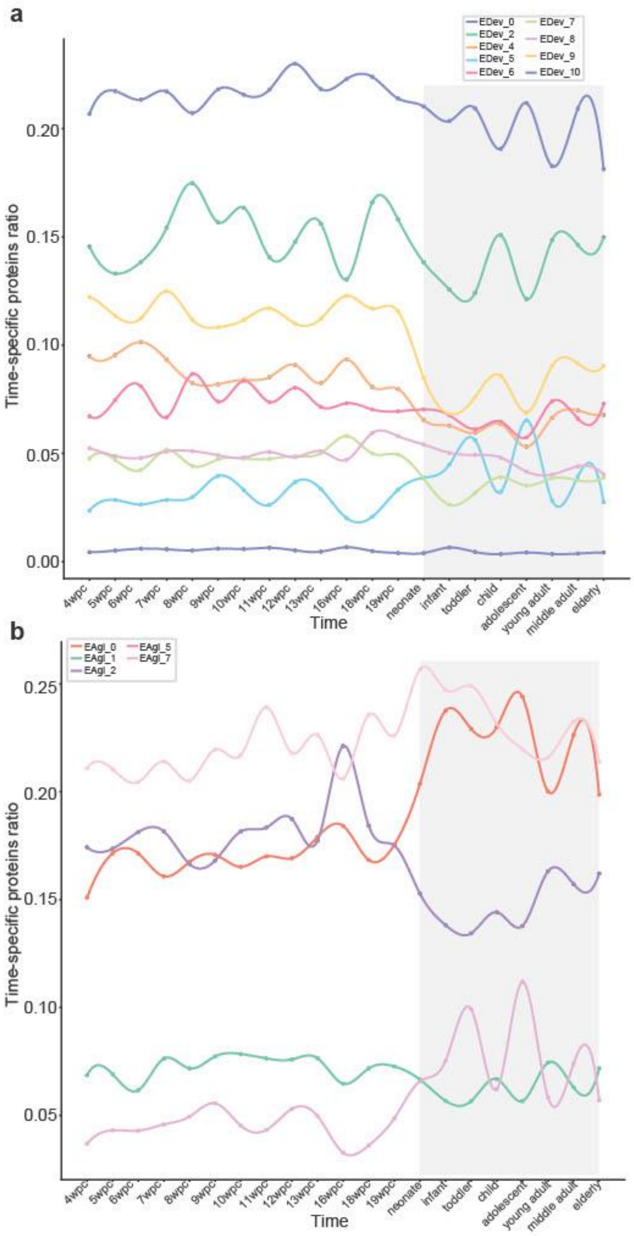


**Supplementary Figure 15**: Proportion of time-specific genes in each Embryological Development (EDev) community (**a**) and Environmental Aging (EAgi) community (**b**) over time. The continuous line was interpolated through a 3-degree spline.

### Supplementary Figure 16

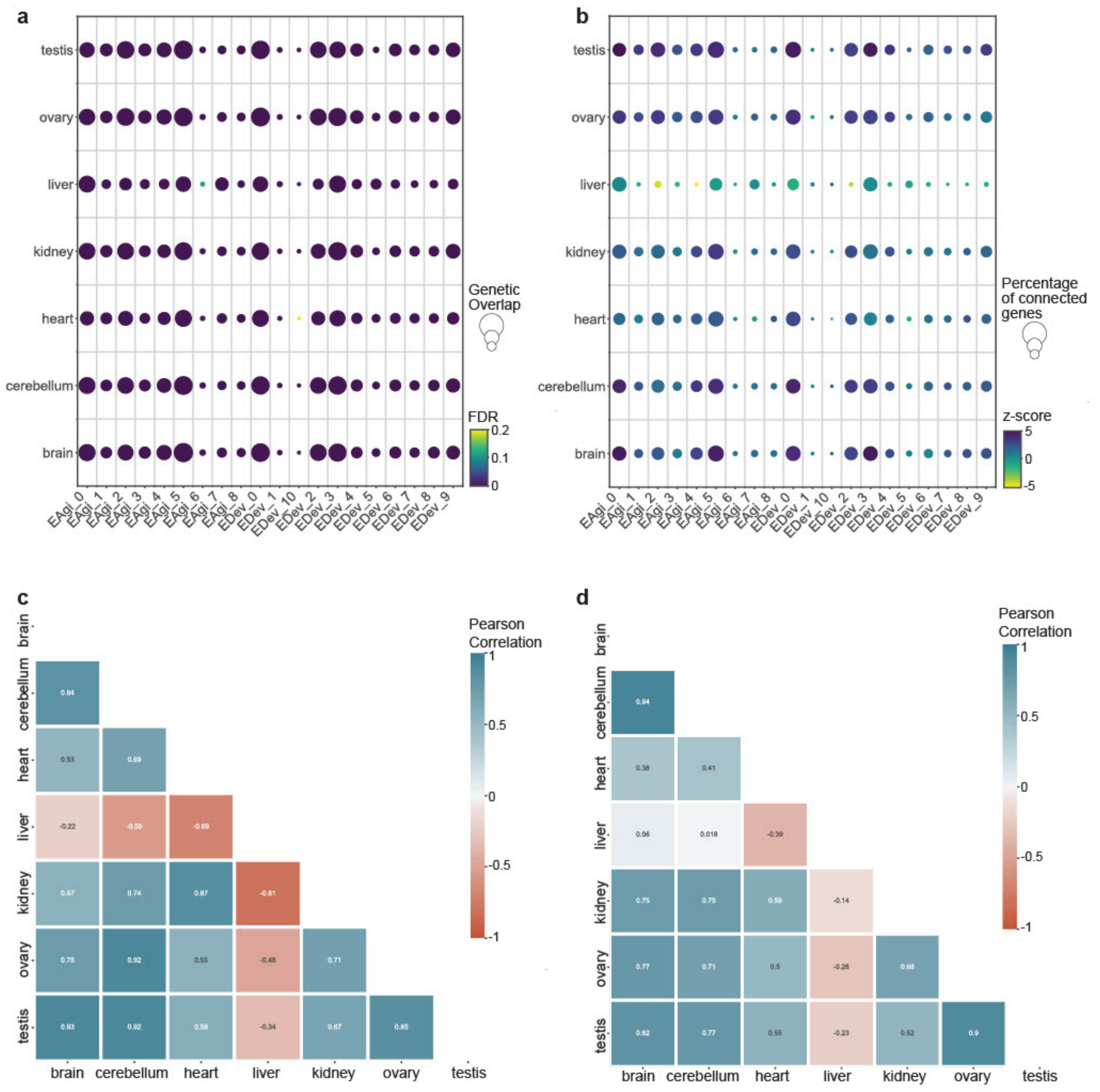


**Supplementary Figure 16**: **a**) Heatmap showing the enrichment of tissue-specific proteins in the EDev and the EAgi communities. The size of each circle is proportional to the number of shared proteins. **b**) Heatmap showing the connectivity z-score of the tissue-specific proteins in the EDev and the EAgi communities. The size of each circle is proportional to the percentage of connected proteins. **c**) Heatmap showing the Pearson's correlation coefficient of the tissue-specific connectivity (lcc) z-score in each EDev community for each tissue pairs. **d**) Heatmap showing the Pearson's correlation coefficient of the tissue-specific connectivity (lcc) z-score in each EAgi community for each tissue pairs.

### Supplementary Figure 17

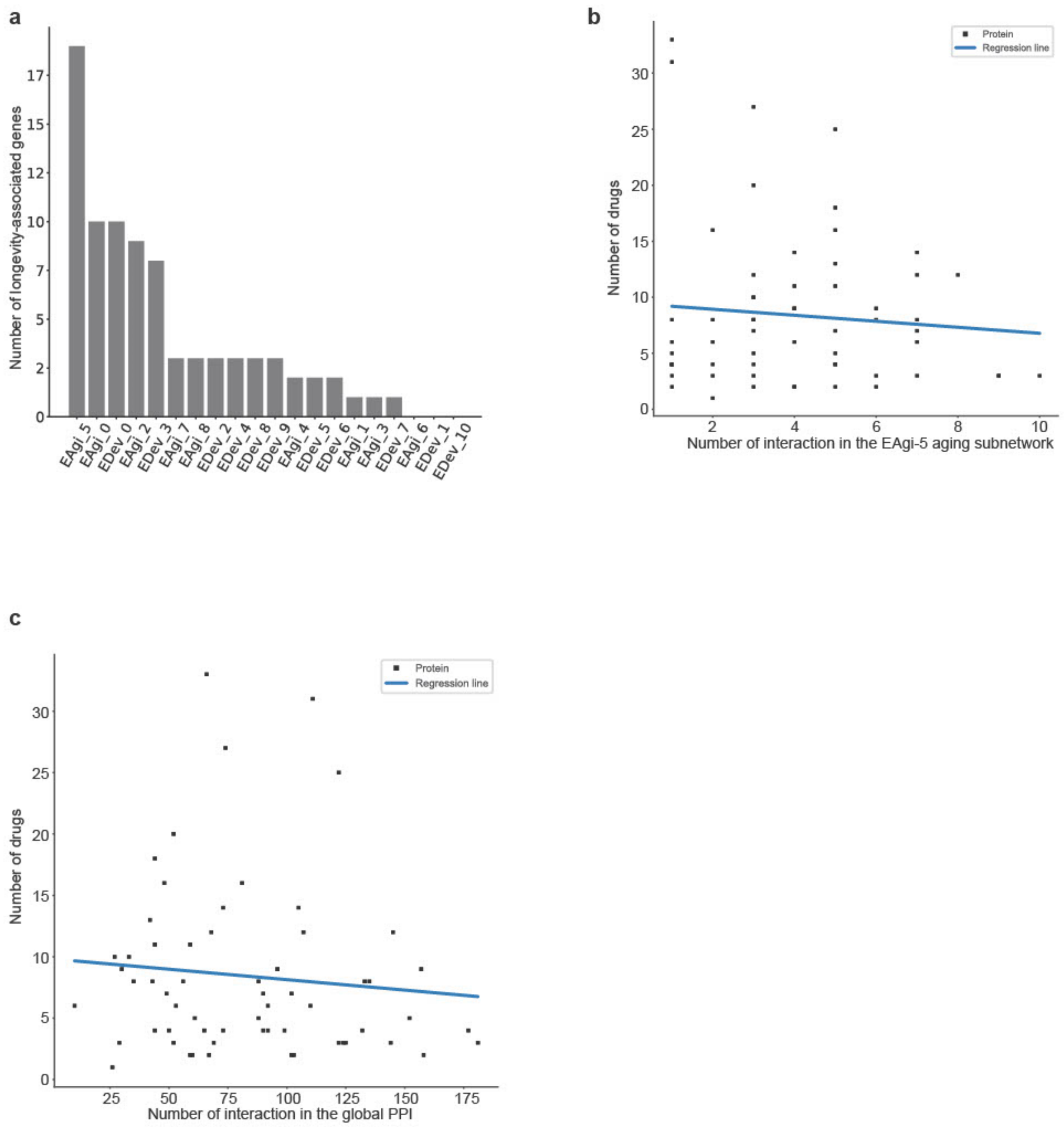


**Supplementary Figure 17**: **a**) Bar plot showing the number of longevity-associated genes associated with each EDev and EAgi community. **b**) Correlation between the number of interactions in the EAgi-5 subnetwork and the number of available drugs for each protein. **c**) Correlation between the number of interactions in the human interactome and the number of available drugs for each protein.
